# Plasmid copy number affects the DNA methylation-driven expression dynamics of the *Cfr*BI restriction-modification system and impacts phage restriction

**DOI:** 10.64898/2025.12.04.692298

**Authors:** Johannes Gibhardt, Trung Duc Hoang, Konstantin Severinov, Mikhail A. Khodorkovskii, Natalia E. Morozova

## Abstract

Restriction-modification (R-M) systems are one of the most widespread and, due to their often plasmid-based nature, transmittable anti-phage systems bacteria have. The *Cfr*BI R-M system studied here consists of a methyltransferase (MT) and a restriction endonuclease (RE) that are divergently expressed and share a promoter region that harbors a single *Cfr*BI recognition site. Previously, the methylation of this site has been shown to regulate the expression of the R-M system. Here, we show that the expression dynamics of the *Cfr*BI R-M system and its protective properties depend on the copy number of the plasmid harboring it. A higher copy number results in higher expression, but the expression on a medium-copy number plasmid interestingly conferred the highest phage resistance. After transformation of naïve cells however, expression of the RE was fastest in the high-copy plasmid background. *In vivo* we show that the expression strength of the MT inhibits its own expression, while enhancing RE expression. To conclude, the results indicate that for phage resistance the overall expression strength might not be the predominant factor in case of the *Cfr*BI R-M system, while for the establishment of the system the initial expression rate of the MT seems to be the determining factor.

## INTRODUCTION

Bacteriophages are one of the oldest known anti-bacterial agents and although they have been a treasure chest for the research of molecular biological principles, their anti-bacterial properties have been neglected for some time. However, due to the emergence of multidrug resistant bacteria and novel discoveries, such as the CRISPR/Cas system, interest in the biology of phages and their interaction with their host bacteria was renewed (1).

In recent years, several approaches have been discussed to utilize phages against bacteria: the use of phages against susceptible bacteria, the delivery of drugs to bacteria, the use of phage-derived proteins, the application of phages in combination with antibiotics to increase the effectiveness and the genetic modification of bacteria with phages to increase antibiotic susceptibility (2). Therefore, phages offer promising approaches to combat multidrug resistant bacteria, but our understanding of the mechanisms bacteria utilize to acquire resistances against phages and how phages overcome them is still limited.

As our understanding of the impact of phages on bacterial ecology, their co-evolution and abundance in nature deepened, new discoveries concerning bacterial resistances against phages were made (1). Besides of the CRISPR/Cas system (3–5), bacteria have evolved a manifold of different systems to counteract infections by phages, such as abortive infection systems, BREX, DISARM, prokaryotic argonaute proteins, Zorya, Theoris and others (6–11). The variety of anti-phage mechanisms demonstrates the close relationship and co-evolution of phages and their bacterial hosts. One of the most widespread, mobile and important systems bacteria utilize against phages, or foreign DNA, are restriction-modification (R-M) systems.

R-M systems consist of a restriction endoribonuclease (RE) that recognizes specific DNA sequences and a methyl transferase (MT) that methylates the same DNA sequence and thereby prevents cleavage by the corresponding RE. This simple organization allows the recognition of self- and the cleavage of foreign DNA, thereby restricting the replication of phage DNA in the bacteria (12).

R-M systems in general can be divided into four types that differ in the number and interaction of the involved proteins, their cofactor and ATP dependencies and their cleavage site in relation to the recognition sequence (12). The often plasmid-based type II R-M systems are generally considered as the most widespread R-M systems with independently acting MTs and REs. In contrast to type I or type III R-M systems, in which protein-protein interaction between MT and RE and (in case of type I) a specificity protein, governs target-site recognition, type II R-M system rely on differential gene expression to regulate methylation or restriction activities (12). Most type II R-M systems consist of the genes encoding a MT and a RE, as well as a control or C protein that is crucial for the regulation of the RE expression (12).

The *Cfr*BI R-M system here studied does not contain a C protein-encoding gene and solely consists of a promoter region and divergently expressed genes encoding for a MT and a RE and thus belongs to the class-II of R-M systems. It was originally isolated and described from the *Citrobacter freundii* 4111 plasmid pZE8 in the 90s as a divergently expressed R-M system with the relaxed recognition sequence of 5’-CCWWGG-3’, with W being A or T nucleotides. The CfrBI RE is therefore an isoschizomer of, among others, *Sty*I, *Eco*130I, *Bss*T1I and *Erh*I (Figure 1A; 13-16). Sequencing results showed that the promoter region between the *cfrBIM* and *cfrBIR* genes, encoding the *Cfr*BI (m^4^C) MT and RE, respectively, contains a unique recognition sequence of the *Cfr*BI R-M system (14). Interestingly, the promoter regions of the divergently expressed *cfrBIM* and *cfrBIR* genes overlap and have been shown to be regulated by N-4C methylation of the *Cfr*BI recognition site that is located in the -35 region of P*_cfrBIM_* and in close proximity to the -10 region of P*_cfrBIR_* (Figure 1A). Methylation interferes with the interaction of σ^70^ of the RNA polymerase and the *cfrBIM* promoter, thereby facilitating binding of the RNA polymerase complex to two less ideal *cfrBIR* promoters, activating their expression (16–18). The expression of the *Cfr*BI R-M system thus is regulated by covalent DNA modification.

**Figure 1.**
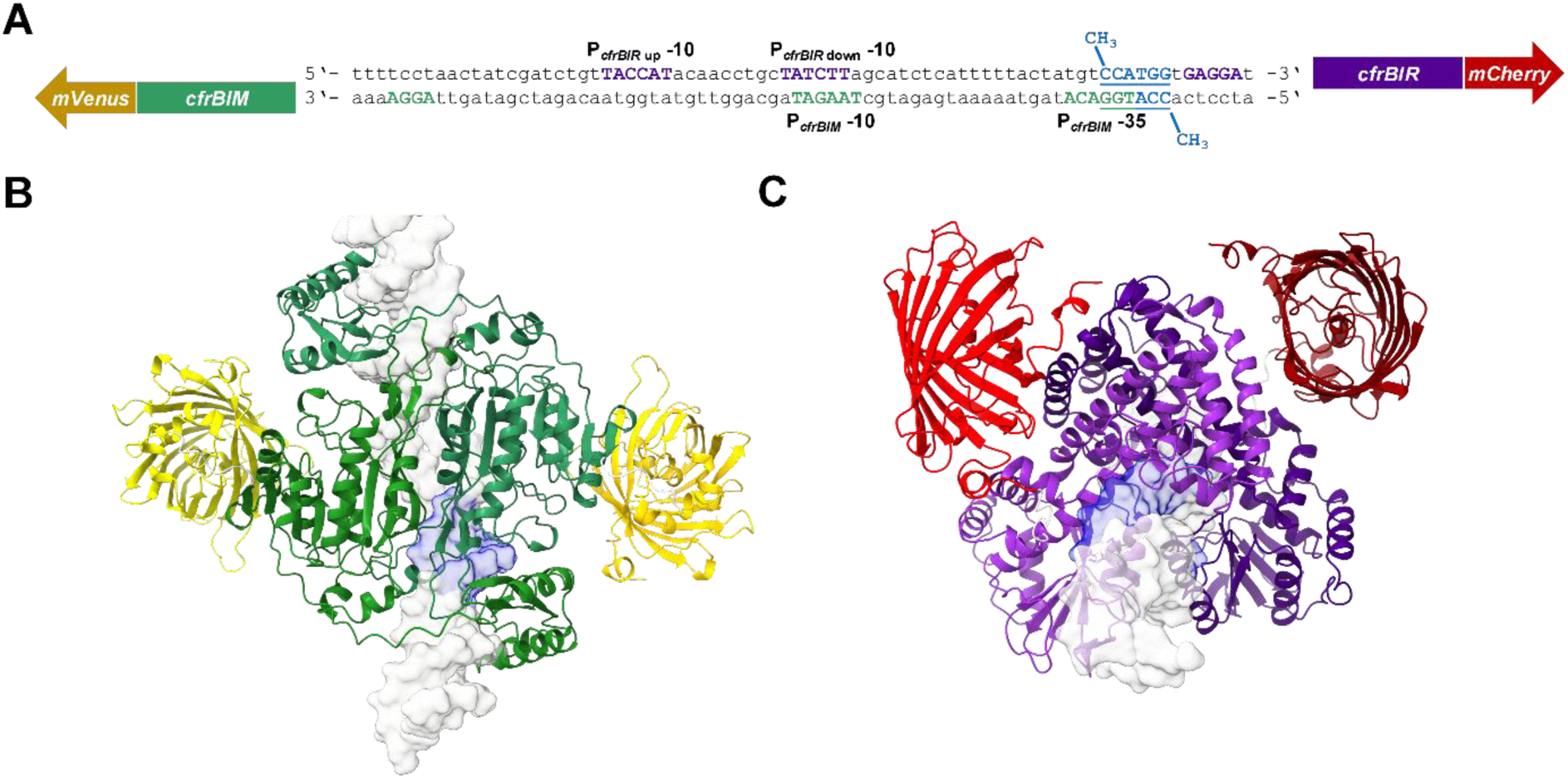
Overview of the *Cfr*BI restriction-modification system. (A) Nucleotide sequence of the intergenic region between the genes *cfrBIM* and *cfrBIR*, encoding the MT and RE of the *Cfr*BI R-M system, respectively. 3’-fusions of *cfrBIM* to *mVenus* and *cfrBIR* to *mCherry* schematically depicted. Important promoter elements depicted in green for the expression of the MT gene and in violet for the RE gene. The intergenic *Cfr*BI recognition sequence that overlaps with the MT promoter and methylation sites are highlighted in blue. (B, C) AlphaFold predictions of the fluorescent fusion proteins, binding to the intergenic region of the *Cfr*BI R-M system. Structures of the MT (B) and RE (C) are depicted in green and violet and the mVenus and mCherry proteins in yellow and red, respectively. The *Cfr*BI recognition sequence is depicted in blue. The models show structural predictions of homodimers. Gene and protein sizes are not drawn to scale.

Since type II R-M systems are often plasmid-based, they are prone to be exchanged via horizontal gene transfer (19). In a naïve cell that is transformed with such a R-M system-containing plasmid it is essential that all available recognition sites of the R-M system are methylated by the MT and therefore no longer substrates for the RE, before the RE is expressed. A failure would result in enzymatic digestion of the chromosome and most likely cell death. Therefore, in a R-M system, such as *Cfr*BI, that is only regulated by the methylation of the *Cfr*BI site in the promoter region of the system, tight regulation of RE expression is instrumental. In the *Escherichia coli* strain DH10B (NC_010473), which is genetically similar to Top10, 1,104 *Cfr*BI recognition sites are present. Furthermore, due to Top10 lacking the *mcrBC* genes, encoding an *E. coli* endonuclease that cleaves DNA containing methylcytosine, expression of the *Cfr*BI MT gene is tolerated by the bacteria (14).

The expression of MT and RE genes in the *Cfr*BI R-M system should strongly depend on the initial expression rate of the MT gene. The faster the available recognition sites are methylated, the faster the expression of the RE gene is activated and the R-M system can fulfil its function, protecting the cells against bacteriophages and foreign DNA. We therefore analyzed the expression dynamics of the *Cfr*BI R-M system the impact of plasmid copy number, and therefore the amount of copies of the *Cfr*BI R-M system, on phage resistance and expression dynamics on the level of bacterial populations and single cells. Our results shed light on the dynamics of this epigenetic regulation of gene expression and how it affects its host ability to protect itself from bacteriophages.

We show that the *Cfr*BI wt R-M system and derivatives containing C-terminal fluorescent protein fusions of the MT and RE are functionally expressed in *E. coli* and protect the bacteria from phage infections. Whether the R-M system is present on high-, medium-, or low-copy plasmids determines the expression dynamics of the MT and RE, as well as the cell growth and the protective properties of the R-M system. Finally, we show that the stronger the MT is expressed, the stronger its own promoter is repressed, which allows for increased expression of the RE.

## MATERIAL AND METHODS

### Bacterial strains, plasmids and growth conditions

*E. coli* Top10 (F^−^*mcr*A Δ(*mrr*-*hsd*RMS-*mcr*BC) φ80*lac*ZΔM15 Δ*lac*X74 *rec*A1 *ara*D139 Δ(*ara-leu*)7697 *gal*U *gal*K λ^−^*rps*L(Str^R^) *end*A1 *nup*G) was used as a host for cloning and all experiments, if not stated otherwise. Oligonucleotides were purchased from Evrogen (Moscow, Russia) or Sigma-Aldrich (Munich, Germany) and are listed in Table 1. *E. coli* strains were grown in lysogeny broth (LB). Agar plates were prepared with 15 g agar/l. *E. coli* transformants were selected on LB plates containing ampicillin (100 µg/ml), tetracycline (10 µg/ml), or chloramphenicol (30 µg/ml). All plasmids used in this study are listed in Table 2. If not otherwise indicated, for experiments with bacteriophages, wt *E. coli* phage λ or the virulent λ_vir_, as indicated, or T7 phages were used and propagated as described earlier (20). Bacteria were grown at 37°C and in liquid cultures shaking at 160 rpm.

**Table 1.**
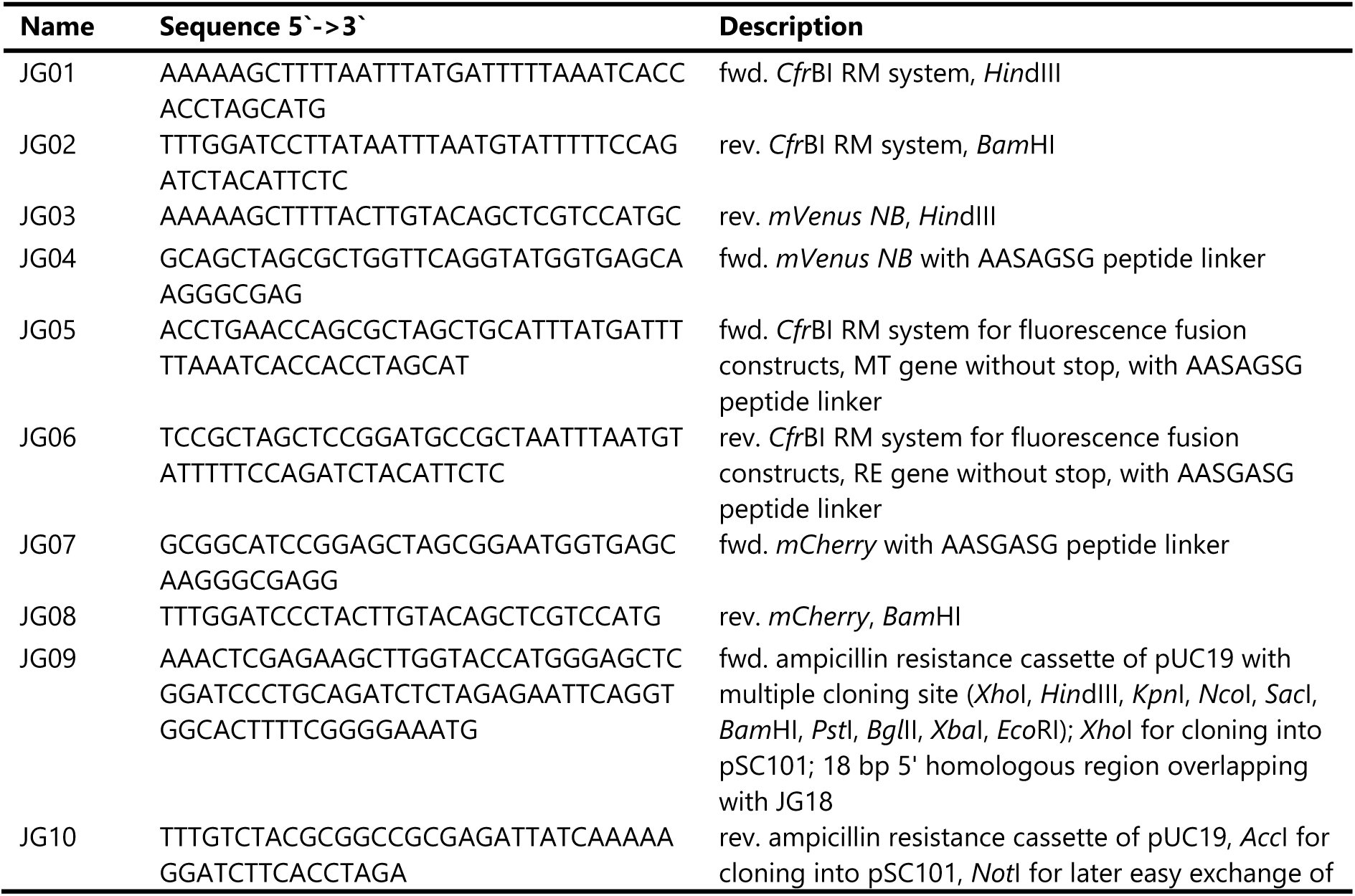
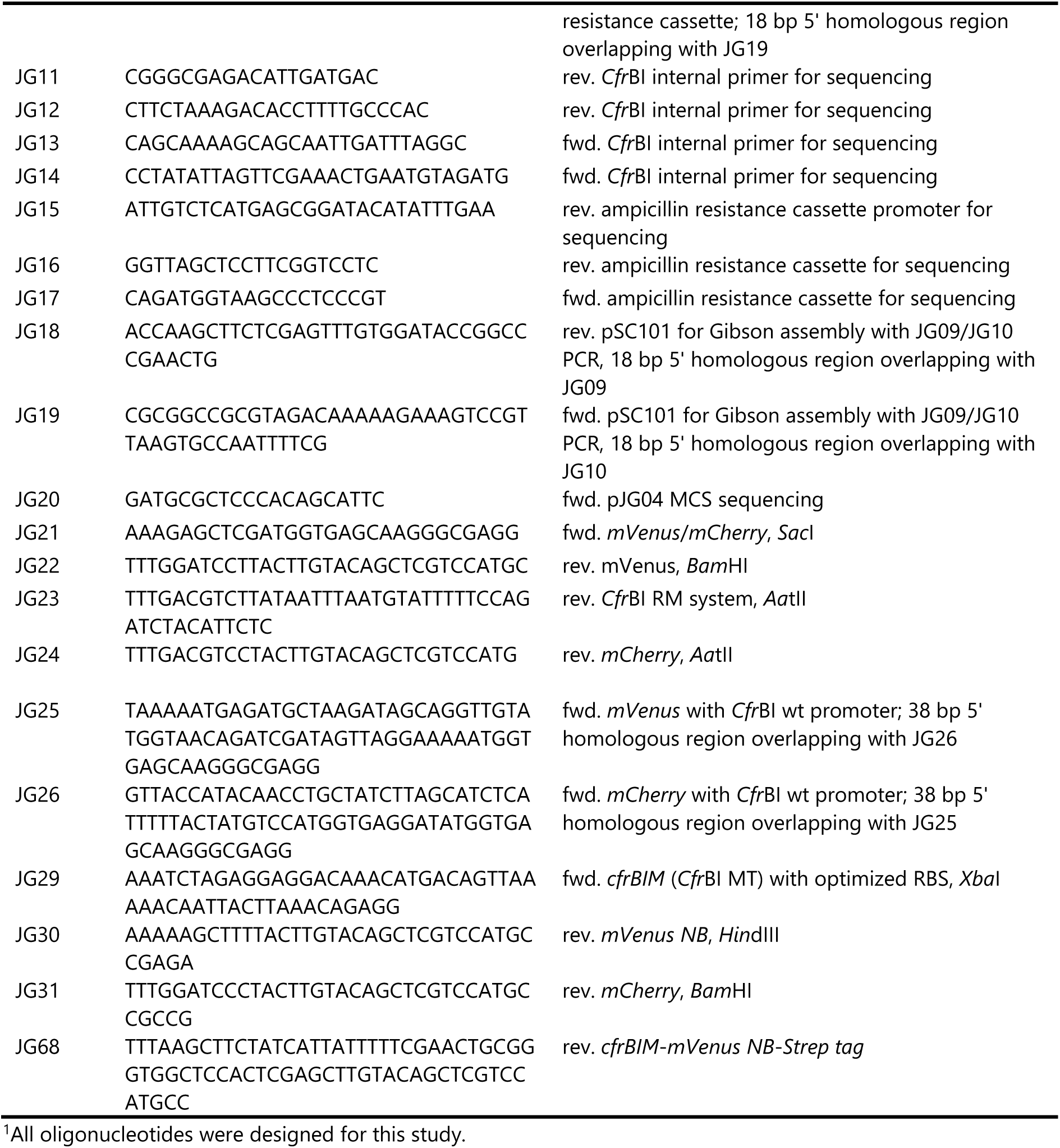
Oligonucleotides.^1^.

**Table 2.**
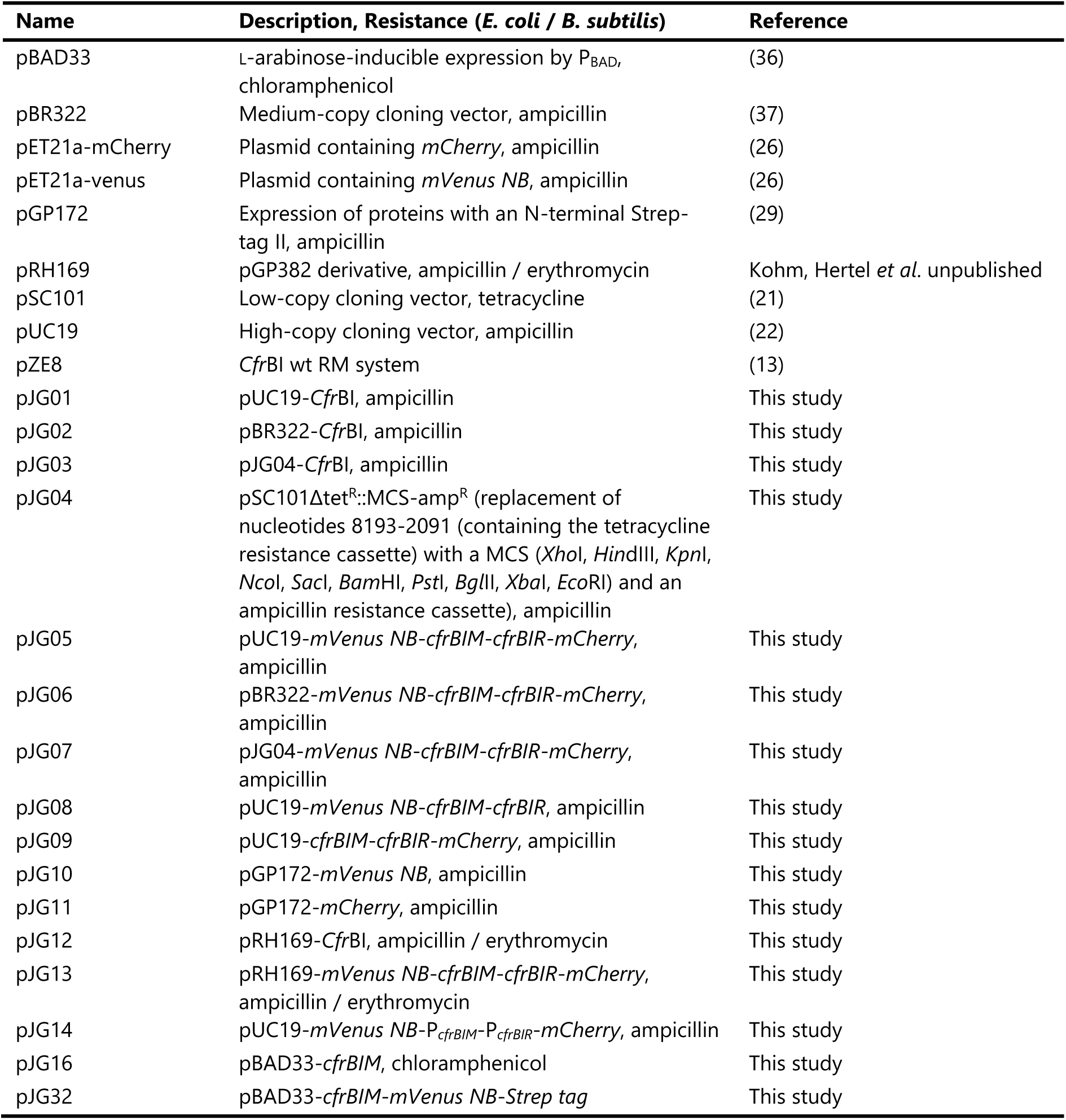
Plasmids.

### DNA manipulation, construction of plasmids and mutant strains

The plasmids that were used and constructed in this study are listed in Table 2. Transformation of *E. coli* was performed using standard procedures (20). Plasmids were isolated from *E. coli* Top10 cultures using the GeneJET Plasmid Miniprep Kit (Thermo Fisher Scientific) or the Monarch Plasmid Miniprep Kit (NEB). PCR products were purified using the GeneJET PCR Purification and Gel Extraction Kits (Thermo Fisher Scientific) and the Monarch PCR & DNA Cleanup and Gel Extraction Kits (NEB). DNA polymerases, restriction enzymes and DNA ligases were purchased from Thermo Fisher Scientific and NEB. Enzymes and Kits were used according to the manufacturer’s instructions. DNA Sanger sequencing was performed by Evrogen (Moscow, Russia) and Microsynth Seqlab (Göttingen, Germany).

The plasmid pJG04 was constructed to have a versatile low-copy vector for the further experiments. Therefore, the backbone of the pSC101 low-copy vector (21) was amplified by PCR using the oligonucleotides JG18 and JG19 and the ampicillin resistance cassette from pUC19 (22) was amplified using the oligonucleotides JG09 and JG10. With JG09 introducing a custom designed multiple cloning site for further cloning applications, consisting of *Xho*I, *Hin*dIII, *Kpn*I, *Nco*I, *Sac*I, *Bam*HI, *Pst*I, *Bgl*II, *Xba*I and *Eco*RI. DNA sequences introduced to the PCR products by the oligonucleotides JG09/JG18 and JG10/JG19, share 18 bp long 5’ homologous regions for Gibson Assembly, respectively. Resulting PCR fragments were joined by Gibson Assembly (23) using the HIFI DNA Assembly Master Mix and the manufactures instructions (NEB), resulting in pJG04, a 7,252 bp derivative of the low-copy pSC101 vector, with the tetracycline resistance cassette containing region (nucleotides 8,193-2,091) replaced by the afore mentioned multiple cloning site and an ampicillin resistance cassette, which if needed can be exchanged using the introduced restriction sties *Eco*RI and *Not*I (see Table 2). Plasmids pJG01, pJG02 and pJG03 were constructed as follows: Using oligonucleotides JG01 and JG02, the region encoding for the *Cfr*BI R-M system (14), was amplified by PCR using the plasmid pZE8 as a template (13). The resulting PCR product was digested with the enzymes *Bam*HI and *Hin*dIII and ligated to *Bam*HI and *Hin*dIII digested pUC19, pBR322 and pJG04, respectively. For plasmids pJG08 and pJG09, the genes encoding for the yellow and red fluorescent proteins mVenus NB (SYFP2, 24) and mCherry (25), were C-terminal fused to the MT and RE genes of the *Cfr*BI system, respectively. Therefore, *mVenus NB* and *mCherry* were amplified by PCR from pET21a-venus and pET21a-mCherry (26) using the oligonucleotide pairs JG03/JG04 and JG07/JG08, respectively. For pJG08, JG03/JG04 amplified *mVenus NB* and JG05/JG02 amplified *Cfr*BI were fused by splicing by overhang extension (SOE) PCR using primer pair JG03/JG12 (27). The resulting PCR product was purified from an agarose gel, *Hin*dIII/*Xho*I digested and ligated to *Hin*dIII/*Xho*I digested and gel-purified pJG01. The resulting plasmid pJG08 expresses the *Cfr*BI MT C-terminal fused to mVenus NB, joined by a short linker sequence adapted from (28) and the wild type *Cfr*BI RE. The pJG09 plasmid, expressing the wild type *Cfr*BI MT and a C-terminal fusion of the RE with a similar short linker sequence (adapted from (28)) and mCherry was constructed in a similar fashion: *mCherry* was PCR amplified using the oligonucleotide pair JG07/JG08, fused to JG01/JG06 amplified *Cfr*BI by SOE PCR (27) using oligonucleotides JG11 and JG08, *Xho*I/*Bam*HI digested, gel-purified and ligated to *Xho*I/*Bam*HI digested pJG01. Finally, plasmid pJG05, containing the C-terminal fusions of *mVenus NB* with the *Cfr*BI MT gene and *mCherry* with the *Cfr*BI RE gene, was constructed by PCR amplification of the *cfrBIR*-*mCherry* fusion of pJG09 using the oligonucleotides JG11/JG08, digestion and subsequent gel purification using the enzymes *Xho*I/*Bam*HI and ligation into *Xho*I/*Bam*HI digested and gel-purified pJG08. For the medium-and low-copy plasmid derivatives pJG06 and pJG07, the whole *Cfr*BI region, including the fusions of MT and RE genes to *mVenus* and *mCherry*, respectively, was amplified from plasmid pJG05 using the oligonucleotides JG03/JG08, digested by *Hin*dIII/*Bam*HI, gel purified and ligated to *Hin*dIII/*Bam*HI digested pBR322 and pJG04, respectively. For the overexpression and purification of mVenus NB and mCherry proteins *via* an N-terminal Strep-tag, plasmids pJG10 and pJG11 were constructed, respectively. The coding sequences were amplified by PCR using pJG08 and pJG09 plasmids, respectively, as templates using oligonucleotide pairs JG21/JG22 for *mVenus NB* and JG21/JG08 for *mCherry* amplification. Resulting PCR products were *Sac*I/*Bam*HI digested and ligated to *Sac*I/*Bam*HI digested pGP172 (29). To assess whether the *Cfr*BI R-M system is able to protect the Gram-positive model bacterium *Bacillus subtilis* from phage infections, plasmids pJG12 and pJG13, expressing the wild type and MT-mVenus-MT/RE-mCherry fusion proteins, respectively, was constructed. The corresponding DNA fragments were amplified using oligonucleotides JG01/JG23 for the wild type and JG03/JG24 for the fusion-protein variants using the plasmids pJG01 and pJG05 as templates. Resulting PCR products were digested using *Hin*dIII and *Aat*II and ligated to the pGP382-based shuttle plasmid pRH169, digested with the same enzymes (Kohm, Hertel *et al*., unpublished) For experiments with *B. subtilis*, strain TS05 (*trpC2*, ΔSPβ, Δ*skin*, ΔPBSX, Δprophage1, *pks*::*cat*, Δprophage 3, *amyE*::P*mtlA*-*comKS*), a derivative of the Δ6 strain TS01 (30) was used (Kohm, Hertel *et al*., unpublished), as described elsewhere (30). To analyze the *in vivo* effect of the methylation of the promoter of the *Cfr*BI MT and RE, plasmids pJG14, expressing *mVenus* from the *Cfr*BI MT promoter and *mCherry* from the *Cfr*BI RE promoters, as well as pJG16, allowing for the L-arabinose-inducible expression of the *Cfr*BI MT, were constructed as follows: *mVenus NB* with the *Cfr*BI promoter region was amplified from pJG10 using oligonucleotides JG25/JG03, *mCherry* with the *Cfr*BI promoter region was amplified from pJG11 using oligonucleotides JG26/JG08 and the resulting PCR products were fused by SOE PCR using oligonucleotides JG30/JG31. The resulting product was gel purified *Hin*dIII/*Bam*HI cut and ligated to pUC19 (*Hin*dIII/*Bam*HI cut). For pJG16, the *cfrBIM* gene, encoding for the MT, was amplified from pJG01 using oligonucleotides JG29/JG01. The resulting PCR product was *Xba*I/*Hin*dIII cut and ligated to pBAD33 (*Xba*I/*Hin*dIII cut). pJG32, for the L-arabinose-inducible expression of the *Cfr*BI MT with a C-terminal fusion to *mVenus NB*-*Strep* tag, was constructed in a similar manner. The genes were amplified from pJG08 using oligonucleotides JG29/JG68. The resulting PCR product was *Xba*I/*Hin*dIII cut and ligated to pBAD33 (*Xba*I/*Hin*dIII cut).

### Infection assays

For infection assays *E. coli* Top10 strains harboring no or the indicated plasmids were adjusted from overnight cultures to an OD_600_ of 0.4 and grown for 1 hour at 37 °C and 160 rpm in λLB. 200 µl of the cultures were mixed with 4 ml λLB, containing 0.4 % (w/v) agar (λLB top agar). This top agar was poured on petri dishes containing 10 ml λLB with 1.5 % (w/v) agar (λLB bottom agar). For *E. coli* Top10 harboring pUC19/pBR322/pJG04 or their derivatives the media contained ampicillin. λLB containing 2·10^9^ λ_vir_ phage particles was 10-fold serial diluted and 5 µl of the indicated dilutions were spotted on the agar plates. Infection assays using T7 phages were performed similar, with the distinction that for quantifications 100 µl cells were mixed with 100 µl phage lysate and top agar, evenly spread across the bottom agar-containing plate. The plates were photographed after 24 hours of incubation at 37 °C. For quantifications, resulting plaques were counted and the plaque forming units per ml phage solution was calculated. Each dilution was spotted 6 times per replicate and strain.

### Growth and fluorescence measurements using a microplate reader

#### Growth experiments of plasmid-containing cells

Overnight cultures of the different strains were used to inoculate LB medium with ampicillin to an OD_600_ of 0.1 and grown at 37 °C with agitation (160 rpm) for 3 to 4 hours. The OD_600_ was adjusted again to 0.1 and 200 µl of the cultures used to inoculate a 96-well plate (Sarstedt). Growth in liquid medium was monitored using a BioTek Synergy H1 microplate reader (Agilent) and 96 well plates (Sarstedt). Data was analyzed using the Gen5 software (v3.16). OD_600_ absorbance was analyzed every 5 or 10 minutes. Fluorescence was detected using an excitation range of 579±15 nm, an emission range of 616±20 nm and a gain of 146 for mCherry. For mVenus NB an excitation range of 509±14 nm, an emission range of 535±20 nm and a gain of 90 was used. For both bottom optics and a read high of 7 mm and xenon lamp flashes with a high lamp energy per data point were used (10 measurements/data point). The bacteria were grown at 37°C with orbital shaking at 237 cpm (4 mm).

#### Growth experiments of naïve cells after transformation

For the analysis of growth and gene expression after transformation of naïve *E. coli* cells, competent Top10 cells were created as described previously (31). To 100 µl frozen aliquots of competent *E. coli* Top10, 100 ng of plasmid DNA were added, mixed and the suspension incubated for 30 minutes on ice. After a 30 second heat shock at 42 °C and subsequent 10 minutes incubation on ice, 500 µl SOB medium (31) were added and the cells grown for 30 minutes at 37 °C and 160 rpm agitation. The OD_600_ was adjusted to 0.1 in 1 ml LB medium, supplemented with ampicillin and 200 µl were used to inoculate a 96-well plate (Sarstedt). The plates were incubated at 37 °C with agitation (237 cpm) and analyzed using the microplate reader with the same settings as described above.

#### Growth experiments of dual plasmid-containing cells

To assess how the methylation of the *Cfr*BI recognition site in the intergenic region between the MT and RE encoding genes affects their expression, *E. coli* cells containing two compatible plasmids were used. Top10 cells harboring either the pUC19 empty vector or a pUC19 derivative with P*_cfrBIM_*-*mVenus NB* and P*_cfrBIR_*_-_*mCherry* fusions (pJG14) was transformed with either pBAD33-*cfrBIM* (pJG16) or the pBAD33 empty vector control. For normalization of the fluorescence signals, a Top10 strain harboring the pUC19 and pBAD33 empty vectors were used. To assess the strength of induction of the MT expression, by the different arabinose concentrations, *E. coli* Top10 harboring the pUC19 empty vector and plasmid pJG32 (pBAD33-*cfrBIM*-*mVenus*-*Strep tag*), which is isogenic to pJG16, was cultivated in parallel. LB overnight cultures of the different strains, supplemented with ampicillin, chloramphenicol and 0.5 % (w/v) D-glucose were used to inoculate LB pre-cultures without glucose that were grown until the exponential growth phase. Finally, the cultures were used to inoculate LB supplemented with ampicillin, chloramphenicol and either 0.5 % (w/v) D-glucose or different L-arabinose concentrations (0.005, 0.001, 0.0005 and 0 % (w/v)) to an OD_600_ of 0.1. 200 µl of the cultures used to inoculate a 96-well plate (Sarstedt) and grown using at 37 °C and the same settings as described above using a microplate reader.

#### Growth experiments in the presence of phages

*E. coli* harboring the different plasmids were inoculated from overnight cultures to an OD_600_ of 0.1, incubated at 37°C and 160 rpm for 30-45 minutes. The OD_600_ was measured again and 200 µl of cells with an OD_600_ of 0.1 with or without addition of wt λ phages corresponding to a multiplicity of infection (MOI) of 0.2 (∼3,2 x 10^6^ PFU) or a MOI of 2.0, (∼3,2 x 10^7^ PFU) were used to inoculate a 96-well plate (Sarstedt) and grown using the same settings as described above using a microplate reader.

### Purification of mVenus and mCherry and normalization of fluorescence units

To normalize the fluorescence units obtained from the microplate reader experiments, calibration curves for mVenus NB and mCherry were created as follows: *E. coli* BL21 harboring pGP172 derivatives for the overexpression of N-terminal *Strep*-tagged mVenus NB and mCherry (pJG10 and pJG11, respectively) were inoculated from overnight cultures to an OD_600_ of 0.1 in 1 l LB medium in 5 l baffled shake flasks and incubated at 37 °C and 160 rpm agitation to an OD_600_ of 0.6 to 0.9. Protein expression was induced by addition of 1 mM IPTG and after an additional 3 hours of incubation the cells were harvested by centrifugation for 15 minutes at 5000 g and 4 °C. The pellets were washed once in ice cold buffer W (0.1 M Tris, 0.15 M NaCl, 0.1 mM Na_2_EDTA, pH 8), centrifuged again for 15 minutes at 3000 g and 4 °C and stored at - 20 °C until further use. Frozen pellets were resuspended in 20 ml ice cold buffer W, supplemented with 1 mM PMSF and lysed using a French press at a pressure of 15000 psi. 1 g diatomaceous earth (Sartorius C100) was added, the mixture vortexed and filtrated using Sartolab RF50 0.22 µm filters (Sartorius). This crude extract was given on 1 ml 50 % *Strep*-Tactin Sepharose resin (IBA, Göttingen), washed 5 times using 1.25 ml buffer W and *Strep*-tagged proteins were finally eluted 5 times using 250 µl buffer E (buffer W with 2.5 mM D-Desthiobiotin. The concentration of the elution fractions was determined using a NanoDrop 2000 (Thermo Fisher) and dilution series were pipette into 96 well plates. Fluorescence was measured using a BioTek SynergyH1microplate reader using the same settings as for the growth experiments (mVenus λ_ex_=509±10nm, λ_em_=535±15nm, gain=90; mCherry λ_ex_=579±10nm; λ_em_=616±20nm, gain=146). The slope was calculated (see Figure 4C; measurements and mean of two replicates) and used for the normalization of the fluorescence units (FU) read outs of the growth curves, normalized to the background fluorescence of the respective empty vector controls.

### Microscopic analysis

Competent *E. coli* Top10 were prepared and transformed with the plasmid pJG05 as described above and immediately analyzed by fluorescence microscopy. 4 µl of freshly transformed cells were given on Agarose pads (1 % agarose in water supplemented with 0.4 % LB medium), on microscopic slides and individual cells subsequently incubated and analyzed by time-laps microscopy. Microscopy was performed using a Nikon Eclipse Ti-E inverted microscope equipped with a custom incubation system. Fluorescence signals for mVenus and mCherry were detected using Semrock filter sets YFP-2427B and TxRed-4040C, respectively. The long-time microscope filming was performed using Micro-Manager 2.0 tools. Time-lapse filming was performed using sCMOS ZYLA 4.2MP Plus Andor camera. Image analysis was performed using ImageJ (Fiji), as described previously (32).

### Structural predictions and data analysis

Structural predictions were performed using the AlphaFold Server (33) website and the default settings. As inputs either wildtype or fluorescent fusion proteins of the *Cfr*BI R-M system and the intergenic region between *cfrBIM* and *cfrBIR* (Figure 1A)were used. Predicted structures were analyzed and rendered using UCSF ChimeraX (University of California, San Francisco;34). For primer design and analysis of sequencing results Geneious Prime (v. 2025.2.2, Docmatics) was used. Microscopic images were analyzed using ImageJ (v. 1.54g; 35). For further data analysis and figure design Office 365 (v. 2509, Microsoft) was used.

## RESULTS

### The *Cfr*BI R-M system and its fluorescent variants confer phage resistance in *E. coli*

The well-known *E. coli* plasmids pUC19 (high-copy plasmid with ∼200 copies per cell; can be as high as 500-700), pBR322 (medium copy plasmid with ∼15-20 copies per cell) and pSC101 (low-copy plasmid with ∼5 copies per cell) were used to elucidate the impact of the copy number on the expression dynamics of the *Cfr*BI R-M system and ultimately phage restriction (20–22, 37–40). To investigate the transcriptional regulation, as well as the ability of the *Cfr*BI R-M system to protect *E. coli* against phages, fluorescent fusion variants were constructed. Since the genes encoding for the MT and RE are transcribed divergently and from a shared promoter region, C-terminal fusions were considered to be less-likely to impact their expression. Protein BLAST using the built in BLASTP suite of REBASE (41) showed that the *Cfr*BI MT and RE share the highest sequence homology with different *Escherichia* and *Salmonella* MTs and REs, respectively. Foldseek blasts, using AlphaFold prediction of monomeric *Cfr*BI MT and dimeric *Cfr*BI RE (filtered for the PDB100 database), revealed highest structural homology of the RE with R.*Bsp*D6I2 (2P14), *Bso*BI RE (1DC1), cA4-activated Card1 (6XL1), Hje (1OB8) and NarL (3EUL) and for the MT with the TTHA0409 MT (2ZIG), a MT of *Adineta vaga* (8S9O), the *Ccr*M MT (6PBD), the *Mbo*IIa MT (1G60), the M1.*Hpy*AVI MT (5HFJ) and the *Pvu*II MT (1BOO).

Interestingly, all of these MTs, with the exception of *Pvu*II, have been crystalized as homodimers. Using AlphaFold Server predictions of the MT and RE proteins with and without C-terminal fluorescent protein fusions together with the intergenic region between *cfrBIM* and *cfrBIR* were modeled. Interestingly, the MT and RE, with or without the C-terminal fusions, are modeled to bind the intergenic region and the *Cfr*BI recognition site as dimers (Figure 1 B & C, supplementary Figure S1). Based on the structural predictions and the genetic context of the *Cfr*BI R-M system, as well as variants that express the MT with a C-terminal mVenus fusion, the RE with a C-terminal mCherry fusion, or both, were cloned into the high-copy *E. coli* vector pUC19 (Figure 2A). Functionality of the *Cfr*BI R-M system in protecting *E. coli* against infections by the *E. coli* phage λ and the ability of the fluorescent protein fusions to complement the wild type phenotype was determined by plaque assays. As shown in Figure 2, *E. coli* cells harboring the empty plasmid are as susceptible to infections by phage λ, as the parental and plasmid-less Top10 strain. In contrast, *E. coli* harboring the wild type *Cfr*BI-R-M system show decreased plaque formation and thus elevated resistance to phage infections. The MT-mVenus, RE-mCherry harboring plasmids, as well as the plasmids harboring both fusions, shows a similar reduction in plaque formation, compared to the wild type R-M system. Thus, the C-terminal fluorescence protein fusions do not impact the ability of the R-M system to restrict phage infections. In the background of the low- and medium-copy vectors pJG04 (pSC101 derivative in which the tetracycline resistance is replaced by an ampicillin resistance gene and a multiple cloning site) and pBR322, respectively, decreased plaque formation can be seen, as well (Figure 3A). The fluorescent protein fusions behave as the wild type variants. Interestingly, the *Cfr*BI R-M system expressed from a low-copy vector shows reduced and from a medium-copy vector elevated phage resistance, compared to the high-copy variant (Figure 3). In case of the *E. coli* phage T7 (supplementary Figure S2), differences between the plasmids harboring the R-M system weren’t as measurable as with phage λ, likely due to the much higher amount of recognition sites (36 in the T7 *vs*. 10 in the λ genome), likely diminishing the concentration of RE needed to efficiently restrict the phage infection.

**Figure 2.**
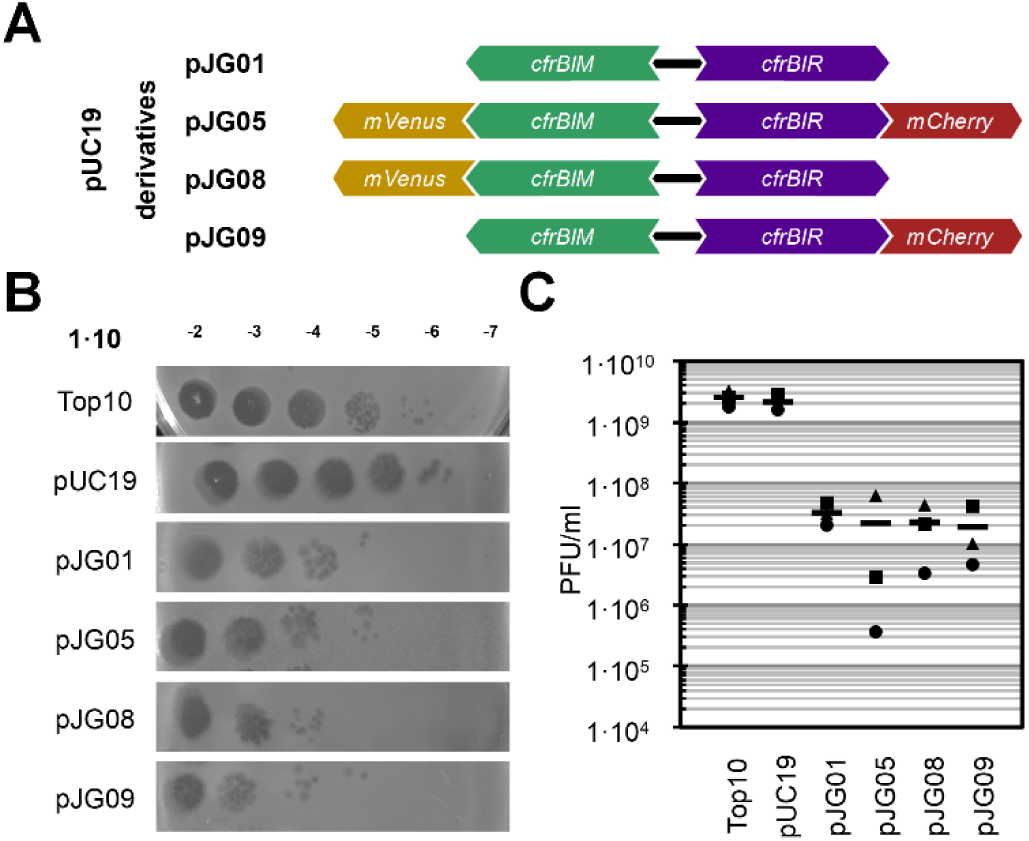
The *Cfr*BI restriction-modification system and its fluorescent protein fusions derivatives confer phage resistance in *E. coli*. (A) Schematic representation of the *Cfr*BI R-M system constructs cloned into the high copy vector pUC19, with pJG01 harboring the wild type *Cfr*BI R-M system, consisting of the divergently expressed genes *cfrBIM* and *cfrBIR* that encode the MT and RE, respectively. The plasmid pJG05 contains the *cfrBIM* gene fused to *mVenus* and the *cfrBIR* gene fused to *mCherry*. The plasmids pJG08 and pJG09 harbor single fusions of *cfrBIM*-*mVenus* and *cfrBIR*-*mCherry*, respectively, while the other gene of the *Cfr*BI R-M system remains in its wild type form. (B) Phage spot assay to assess phage restriction by the different *Cfr*BI R-M constructs using *E. coli* Top10 strains harboring no or the indicated plasmids (see (A)). λLB containing 2·10^9^ λ_vir_ phage particles was 10-fold serial diluted and 5 µl of the indicated dilutions were spotted on agar plates overlaid with λLB top agar, containing the different *E. coli* strains. The plates were photographed after 24 hours of incubation at 37 °C. (C) Graphic representation of the estimated plaque forming units. Plaques of three biological replicates (circle, triangle and square; as described in (B)) of *E. coli* Top10 strains harboring no or the indicated plasmids (see (A)) were counted and the plaque forming units per ml phage solution was calculated. Each dilution was spotted 6 times per replicate and strain. The bars represent the mean value of the three replicates.

**Figure 3.**
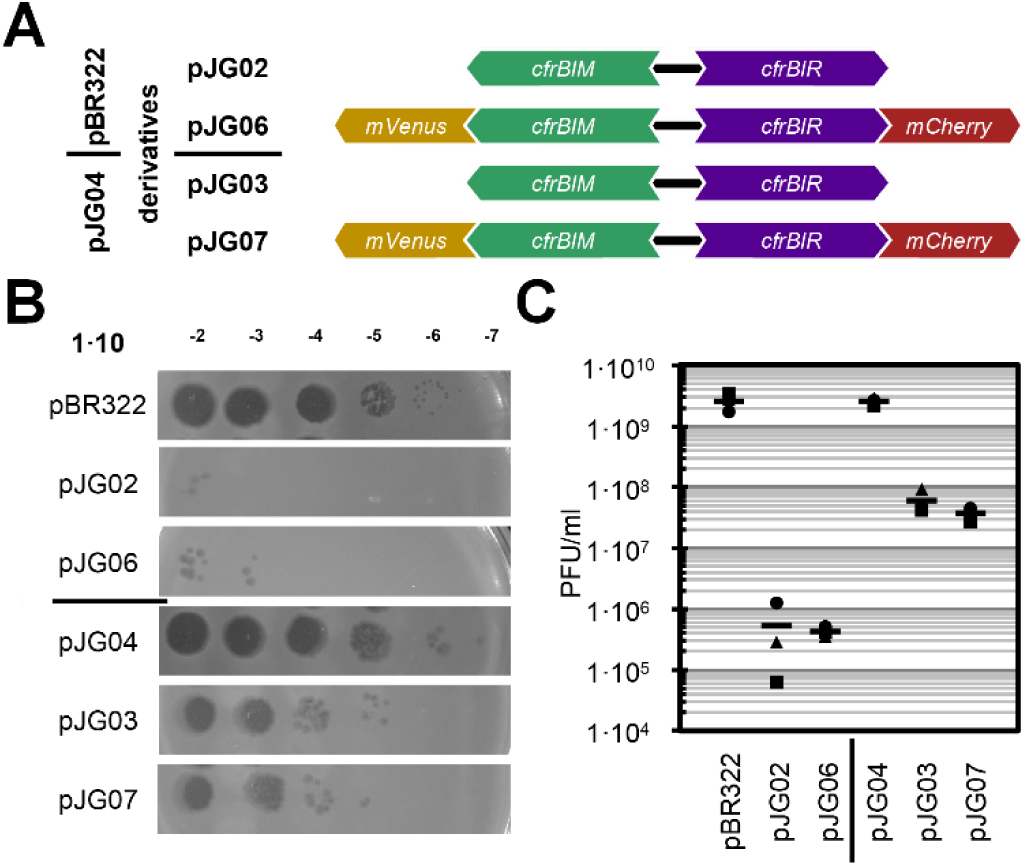
Plasmid copy number affects phage resistance in *E. coli*. (A) Schematic representation of the *Cfr*BI R-M system constructs cloned into the medium copy vector pBR322 (pJG02 and pJG06) or the low copy vector pJG04 (pJG03 and pJG07), which is based on the pSC101 plasmid. pJG02 and pJG03 harbor the wild type *Cfr*BI R-M system, consisting of the divergently expressed genes *cfrBIM* and *cfrBIR* that encode the MT and RE, respectively and plasmids pJG06 and pJG07 contain the *cfrBIM* gene fused to *mVenus* and the *cfrBIR* gene fused to *mCherry*. (B) Phage spot assay to assess phage restriction by the different *Cfr*BI R-M constructs using *E. coli* Top10 strains harboring the indicated plasmids (see (A)). λLB containing 2·10^9^ λ_vir_ phage particles was 10-fold serial diluted and 5 µl of the indicated dilutions were spotted on agar plates overlaid with λLB top agar, containing the different *E. coli* strains. The plates were photographed after 24 hours of incubation at 37 °C. (C) Graphic representation of the estimated plaque forming units. Plaques of three biological replicates (circle, triangle and square; as described in (B)) of *E. coli* Top10 strains harboring the indicated plasmids (see (A)) were counted and the plaque forming units per ml phage solution was calculated. Each dilution was spotted 6 times per replicate and strain. The bars represent the mean value of the three replicates.

These results (Figure 2 & 3) show that the *Cfr*BI R-M system confers phage resistance to *E. coli*, that the fluorescent protein fusions complement the function of the wild type variants and that the effectiveness of the R-M system in preventing infections by phage λ varies with the copy number of the plasmids harboring it.

### Expression dynamics are plasmid-dependent

Next, we investigated the expression dynamics of the MT and RE of the *Cfr*BI R-M system using the fluorescence fusion proteins and how the plasmids from which they are expressed impact these. As seen in Figure 4, growth of the strains expressing the fluorescence fusion proteins did not deviate from the corresponding empty vector controls in the case of the medium- and low-copy plasmids. However, strains harboring high-copy plasmids grew slower in general and those expressing the R-M system showed further decrease in growth that, although not severe, is replicable. *E. coli* harboring the pUC19-derived plasmids must endure higher metabolic burden and higher expression of the plasmid-encoded genes, leading to a reduced growth rate, compared to the medium- or low-copy plasmid-containing cells.

**Figure 4.**
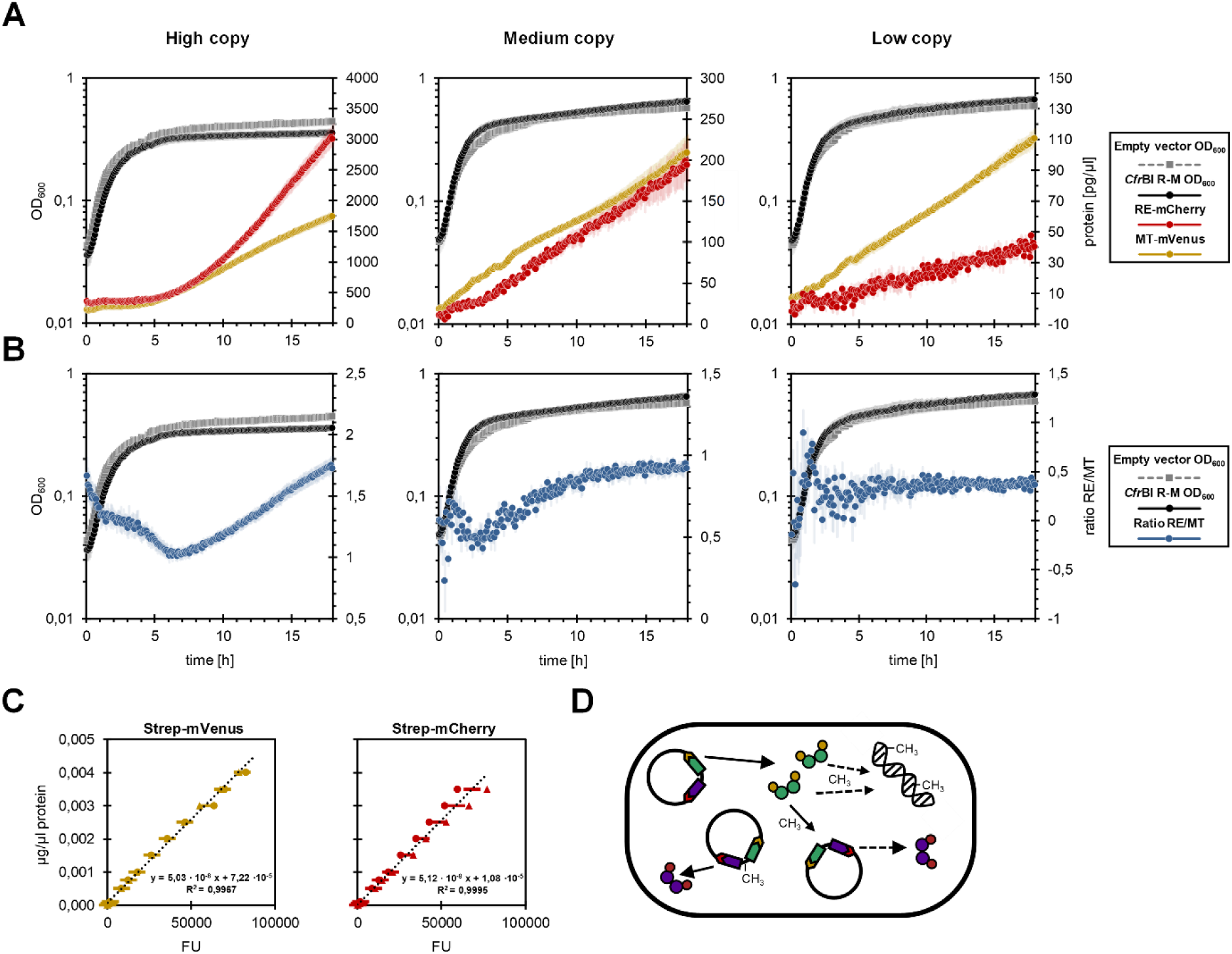
Plasmid copy number affects the expression dynamics of the *Cfr*BI restriction-modification system in *E. coli*. (A) *E. coli* Top10 harboring the plasmids pJG05, pJG06, or pJG07, or the respective empty vector controls (pUC19, pBR322 and pJG04), were analyzed using a microplate reader. 200 µl of cultures per well with an OD_600_ of 0.1 in LB medium containing ampicillin were incubated at 37 °C with agitation (237 cpm) for 18 hours and the OD_600_ and fluorescence were measured every five minutes. The protein concentrations were calculated from the fluorescence units using the slopes of the calibration curves (see C) and normalized to the background fluorescence of the corresponding empty vector control. Shown are the growth of *E. coli* harboring the depicted plasmids or their corresponding empty vector controls, as well as the changes in MT-mVenus and RE-mCherry concentrations in cultures of *E. coli* harboring the depicted plasmids. (B) Growth (OD_600_) of *E. coli* harboring the depicted plasmids or their corresponding empty vector controls (see A) and the ratio of RE-mCherry to MT-mVenus expression, calculated from their concentrations in (A), are shown. (C) N-terminal Strep-tagged mVenus and mCherry proteins were expressed and purified (see material and methods). The concentration of the elution fractions was determined, and the fluorescence of dilution series were measured using the same settings as for the growth experiments. The slope was calculated and used for the normalization of the fluorescence units (FU) read outs of the growth curves (shown are the means of two replicates). (D) Schematic overview of the MT-mVenus and RE-mCherry expression and their action during the experiment. Data represents the mean of four replicates and their standard error (half-transparent shades).

The total strength of expression of MT-mVenus and RE-mCherry corresponds with the plasmid copy numbers, with the high-copy plasmids resulting in the highest overall expression. To account for differences between the mVenus and mCherry signals, the arbitrary fluorescence units were normalized using dilutions of *Strep*-tag purified mVenus and mCherry proteins, measured using the same settings (Figure 4C). Interestingly, not only the pure strength of expression is impacted by the plasmid backgrounds, but also their dynamics. In the high-copy background, expression of the RE-mCherry exceeds the expression of the MT-mCherry and accumulates in later growth phases. For the medium-copy plasmid, expression ratios are quite similar, while in the low-copy background, expression of the RE-mCherry is weaker compared to the MT-mVenus throughout the experiment. Comparing the ratios of MT/RE expression (Fig 4B), for the high- and low-copy plasmids, the ratio rises until it switches and begins to decrease. In the background of the low-copy plasmid, the signal to noise ratio at the beginning is too low and in later growth phases a stabilization is seen.

In the experimental setups used to assess phage restriction and expression dynamics of the *Cfr*BI R-M system, freshly diluted cells, after short precultures were either exposed to phages or their growth was assessed (Figure 2-4). Looking at the expression dynamics of the MT and RE genes, in all three cases, expression of the MT increases, before expression of the RE (to different extends) resumes.

The high accumulation of the RE in cells containing the high-copy plasmids might interfere with the epigenetic switch. Although hemimethylation of the recognition site is sufficient to block cleavage by restriction endonucleases (42), hemimethylation of the cytosine overlapping with the -35 element of the MT promoter is sufficient to alter the expression of MT and RE (18). Highly overexpressed RE, as in the case of the high-copy plasmids might interfere with the binding of the MT to their shared recognition site, thus explaining why the expression patterns towards the RE change later in cells harboring the high- *vs*. the low-copy plasmid, despite more copies favoring stronger MT expression.

Since R-M systems are, due to their often plasmid-based nature, prone to be distributed via horizontal gene transfer, we further investigated the expression of MT-mVenus and RE-mCherry after transformation of naïve cells with the *Cfr*BI R-M system in the background of the different plasmids (Figure 5). At the beginning of the growth curve, OD_600_ consists of non-transformed cells and transformed cells. Since all plasmids confer an ampicillin resistance as a selective marker, the successfully transformed cells afterwards quickly outcompete their non-resistant competitors. Similar to cells already harboring the different plasmids, expression of the MT-mVenus and RE-mCherry again correlated with the plasmid background (high>medium>low). Interestingly, when looking at the ratios of RE to MT expression, at the beginning signal to noise ratio is too low, but in all three cases a stabilization is visible with the high-copy plasmids having detectable MT and subsequent RE expression after 3-4 hours, medium-copy plasmid-containing cells after 4-5 hours and low-copy plasmids only after 7-8 hours after transformation. This is in agreement with the initial hypothesis that faster and higher expression of the MT favors a faster switch to RE expression. Expression of the *Cfr*BI R-M system from higher copy plasmids thus may favor restriction of phage restriction shortly after transformation, but after its establishment the more balanced expression, without the overexpression of the proteins that seems to negatively impact the expression dynamics (Figure 4), from medium copy plasmids seems to be more beneficial and less costly for their host. Also, on the level of single cells (Figure 5C), expression of the MT precedes the RE expression. Naïve *E. coli* cells that have newly taken up the R-M system-containing plasmids show observable expression of the MT gene about 45 minutes after transformation, while expression of the RE lags for another 90 minutes.

**Figure 5.**
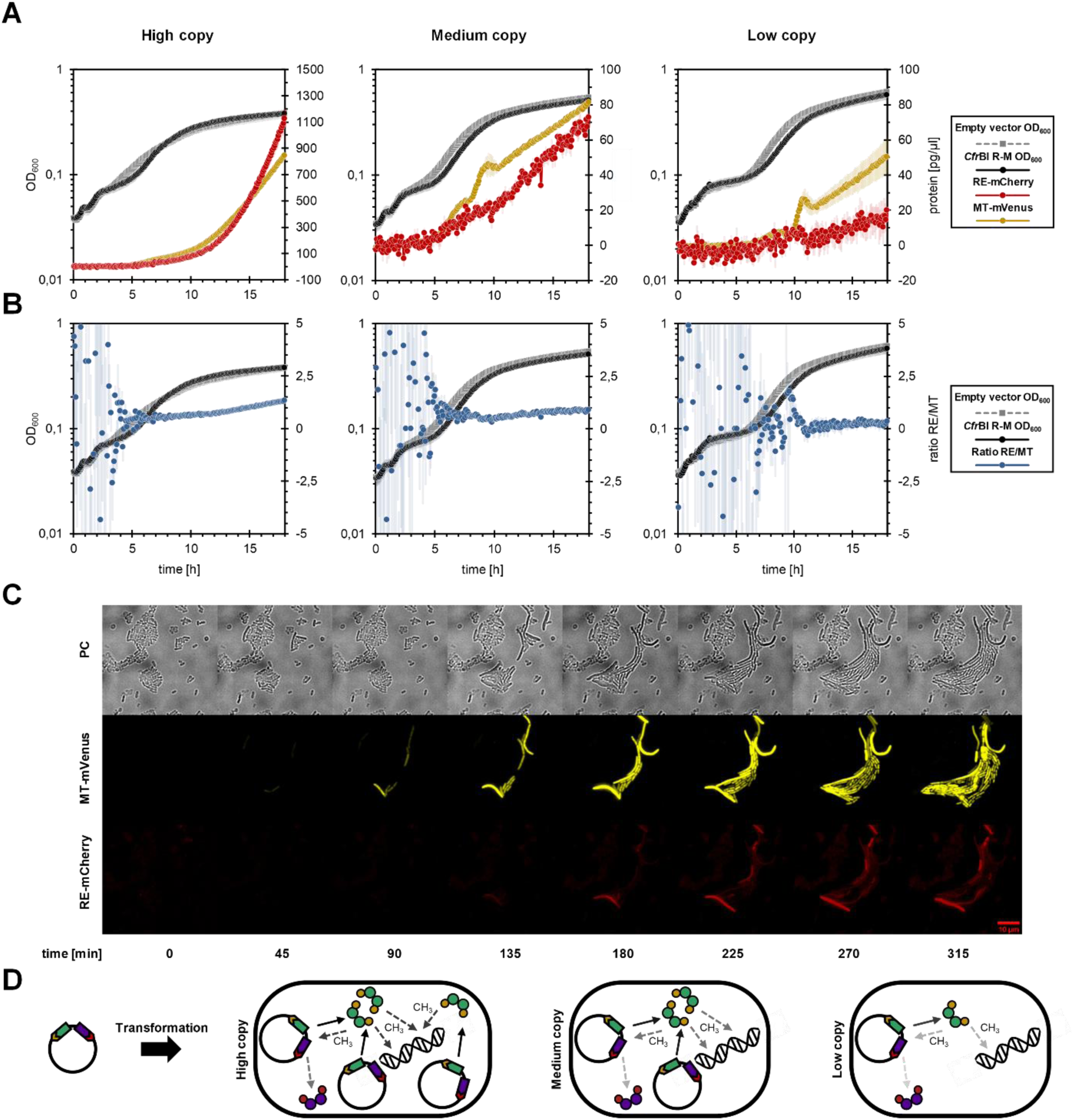
Plasmid copy number affects the expression dynamics of the *Cfr*BI restriction-modification system after transformation of naïve *E. coli* cells. (A) Competent *E. coli* Top10 were transformed with the plasmids pJG05, pJG06, or pJG07, or the respective empty vector controls (pUC19, pBR322 and pJG04) and immediately analyzed using a microplate reader. 200 µl of cultures per well with an OD_600_ of 0.1 in LB medium containing ampicillin were incubated at 37 °C with agitation (237 cpm) for 18 hours and the OD_600_ and fluorescence were measured every five minutes. The protein concentrations were calculated from the fluorescence units using the slopes of the calibration curves (see Figure 4C) and normalized to the background fluorescence of the corresponding empty vector control. Shown are the growth of *E. coli* transformed with the depicted plasmids or their corresponding empty vector controls, as well as the changes in MT-mVenus and RE-mCherry concentrations in cultures of *E. coli* transformed with the depicted plasmids. (B) Growth (OD_600_) of *E. coli* after transformation with the depicted plasmids or their corresponding empty vector controls (see A) and the ratio of RE-mCherry to MT-mVenus expression, calculated from their concentrations in (A), are shown (zoomed in while neglecting a few points during the initial noise). (C) *E. coli* cells transformed with pJG05 were microscopically analyzed on Agarose pads (supplemented with LB medium), using a Nikon Eclipse Ti-E inverted microscope (see material and methods). PC=phase contrast. (D) Schematic overview of the MT-mVenus and RE-mCherry expression and their action after transformation of naïve *E. coli* cells. Growth data represents the mean of four replicates and their standard error (half-transparent shades).

To conclude, expression strength, dynamics, as well as the dynamic ratio between RE to MT expression depends on the plasmids harboring the *Cfr*BI R-M system. The higher the copy number, the higher the overall expression and ultimately higher RE to MT ratio. In the background of the medium-copy plasmid, the switch from MT to RE expression occurs earlier than in the background of the high-copy plasmid. At the same time, growth of cells containing the high-copy plasmid is slower due to increased metabolic burden, which the plasmid replication and high expression of the R-M system impose on the cells (Figure 4). After transformation of naïve cells, expression of MT and subsequently RE commences earlier with increasing copy number of the plasmid harboring the R-M system (Figure 5).

### Regulation of the MT and RE promoters by DNA-methylation

As has been described before, methylation of the cytosine of the *Cfr*BI recognition site that overlaps with the -35 element of the MT promoter, inhibits expression of the MT and thereby facilitates expression of the two weaker RE promoters (16, 18). To further investigate how the DNA-methylation of the promoter region between MT and RE encoding genes regulated their expression and especially the dynamics of this regulation, a dual vector system was used. *E. coli* cells were transformed with two compatible plasmids, the L-arabinose-inducible pBAD33 (36), from which the *Cfr*BI MT was expressed and a pUC19 derivative harboring the promoter region of the *Cfr*BI R-M system, with the open reading frames of the MT and RE encoding genes replaced by *mVenus* and *mCherry*, respectively. First, we wanted to assess the strength of MT expression under the settings used. Therefore, a strain harboring the pUC19 empty vector and a pBAD33 that expresses the *Cfr*BI MT fused C-terminal to mVenus NB-*Strep* tag was used. As shown in Figure 6A, the strength of expression of the MT-mVenus fusion protein increases with increased L-arabinose addition. Next, the expression of mVenus (driven by the MT promoter) and mCherry (driven by the RE promoters) and the impact of the non-fluorescing *Cfr*BI wt MT, using the same L-arabinose concentrations as in Figure 6A, was investigated. Therefore, an *E. coli* strain harboring the reporter plasmid pJG14 (pUC19-*mVenus NB*-P*_cfrBIM_*-P*_cfrBIR_*-*mCherry*) and the plasmid pJG16 (pBAD33-*cfrBIM*) was employed. As shown in Figure 6B, at low concentrations of L-arabinose the activity of both promoters can be detected, with the MT promoter-driven expression being the strongest. With increasing concentrations of L-arabinose and therefore MT expression however, expression of the MT promoter ceases nearly completely, while expression of the RE increases nearly 2-fold. An isogenic strain with the pBAD33 empty vector, grown under the same conditions did not show any alterations in RE or MT expression. Independent of the concentration used, growth and fluorescence were comparable to the data shown in Figure 6B without the addition of L-arabinose (supplementary Figure S3). Higher concentrations of arabinose did not affect expression patterns any further (data not shown). The observed expression patterns of the promoter-reporter gene fusions in the background of the pUC19 plasmid differ from the natural expression pattern of pUC19 harboring the full *Cfr*BI R-M system. This is likely due to the lack of negative auto regulation of the MT expression (16, 18).

**Figure 6.**
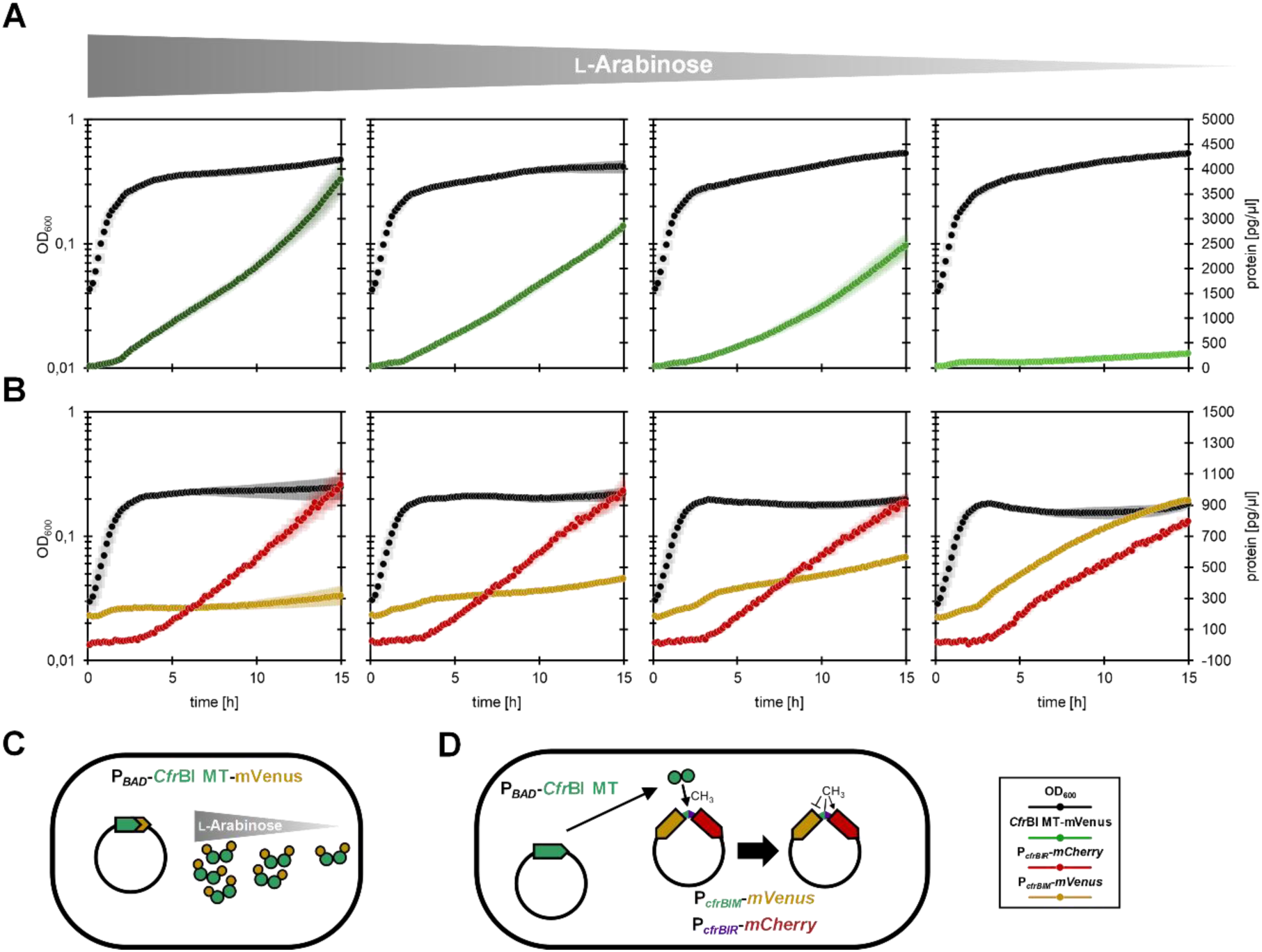
Strength of methyltransferase expression affects the expression dynamics of the *Cfr*BI restriction-modification system. (A) *E. coli* Top10 harboring pJG32 (pBAD33-*cfrBIM*-*mVenus*-*Strep* tag) pJG16 (pBAD33-*cfrBIM*), expressing the *Cfr*BI methyl transferase (MT) with a C-terminal mVenus-Strep tag fusion, and the pUC19 empty vector were analyzed using a microplate reader. 200 µl of cultures per well with an OD_600_ of 0.1 in LB containing ampicillin, chloramphenicol and different _L_-arabinose concentrations (from left to right: 0.005, 0.001, 0.0005 and 0 % (w/v)) were incubated at 37 °C with agitation (237 cpm) for 15 hours and the OD_600_ and fluorescence were measured every 10 minutes. The protein concentrations were calculated from the fluorescence units using the slopes of the calibration curves (see Figure 4C) and normalized to the background fluorescence of the corresponding empty vector control. Shown are the growth of *E. coli*, as well as the changes in MT-mVenus concentrations. (B) *E. coli* Top10 harboring pJG16 (pBAD33-*cfrBIM*), expressing the *Cfr*BI methyl transferase, and pJG14, a pUC19-derivative that contains P*_cfrBIM_*-*mVenus* and P*_cfrBIR_*-*mCherry* fusions were analyzed using a microplate reader, as described in (A). Shown are the growth of *E. coli*, as well as the changes in mVenus and mCherry expression, driven by P*_cfrBIM_* and P*_cfrBIR_*, respectively. (C) Schematic overview of the MT-mVenus-Strep tag expression for the experiment shown in (A). (D) Schematic overview of the MT expression and its impact on the expression of the *cfrBIM* and *cfrBIR* promoters for the experiment shown in (B). Data represents the mean of three replicates and their standard error (half-transparent shades).

The pBAD33 plasmid harbors exactly one *Cfr*BI recognition site in the *cat* gene. Therefore, methylation by the MT does not impact the expression of the P*_BAD_* promoter. Comparing the expression dynamics, as seen in the for the full R-M system (Figure 4), with those of the dual-reporter system (Figure 6), the consequences of the autoregulation of MT expression become obvious with the total inhibition of the MT promoter in the latter.

Since R-M systems are often described as prone to be distributed by horizontal gene transfer, we also evaluated whether the *Cfr*BI RM system is functional in the Gram-positive model bacterium *B. subtilis*. However, neither expression of the fluorescence fusion variant, nor restriction of phage infections by the *B. subtilis* phage SPβ could be detected (data not shown). Furthermore, using the dual plasmid approach, no expression from the wild type *Cfr*BI R-M promoters, nor with an optimized ribosomal binding site could be detected (data not shown), indicating that the promoters are non-functional in *B. subtilis*.

To conclude, using a dual-reporter system we demonstrate the dynamic effects of MT expression on the promoters of the *Cfr*BI R-M system *in vivo*. Increased MT expression directly results in a decrease of the MT promoter activity, while increasing the activity of the two RE promoters (Figure 6). Furthermore, the transfer of the *Cfr*BI R-M system by horizontal gene transfer seems to be limited to proteobacteria.

### Infection dynamics

Finally, we wanted to assess how the dynamic growth behavior and expression of the MT and RE genes are influenced by the presence of phages. Therefore, we cultivated *E. coli* harboring the high, medium- and low-copy plasmids, with or without the fluorescing *Cfr*BI R-M system, in the absence or presence of the *E. coli* phage λ (Figure 7). In the absence of phages (Figure 7A), growth and expression of the MT and RE genes was similar to the previous observations (Figure 4). The presence of phages, independent of their concentration, did not affect expression of the MT or RE encoding genes and growth of the cells harboring the *Cfr*BI R-M system (Figure 7B & C). However, growth of the *E. coli* strains harboring the corresponding empty vector controls was affected differently. In the case of the medium- and low-copy plasmids, a slight inhibition and later recovery of growth is visible at a lower phage to cell ratio (MOI of 0.2), while the phenotype becomes more pronounced at a higher phage to cell ratio (MOI of 2.0). In contrast, the *E. coli* cells harboring the high-copy pUC19 empty vector are nearly as affected at a MOI of 0.2 as they are at a MOI of 2.0. Interestingly, similar to the growth phenotypes observed earlier (Figure 4), *E. coli* harboring the high-copy plasmids show slower growth, compared to those harboring medium- or low-copy plasmids, corroborating earlier observations. The increased metabolic burden the maintenance of high-copy plasmids and increased gene expression means for the host cells thus increases the susceptibility for phage infections (Figure 7).

**Figure 7.**
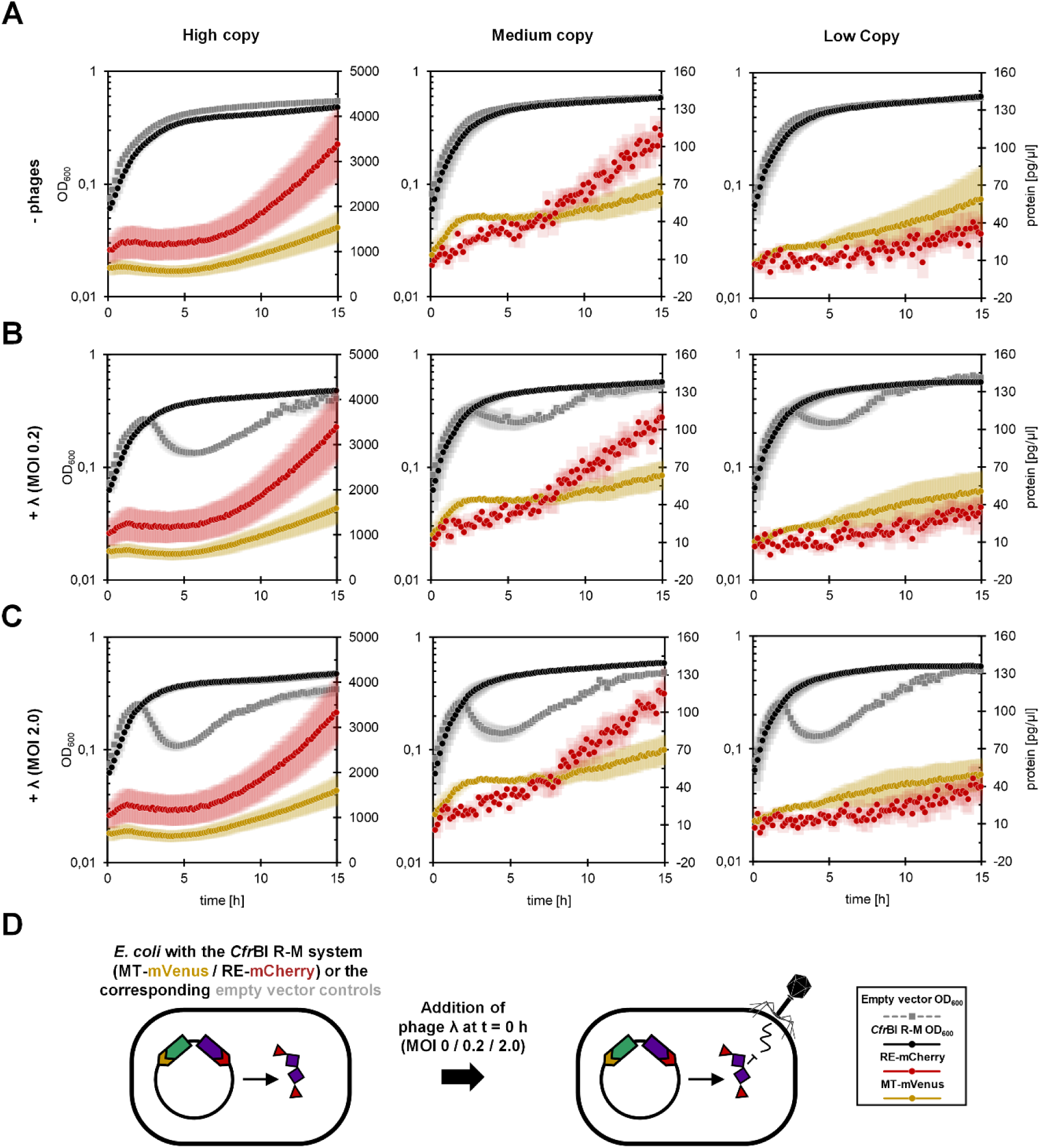
Plasmid copy numbers affect the susceptibility of *E. coli* to phage infections in dynamic infection assays. *E. coli* Top10 harboring the plasmids pJG05, pJG06, or pJG07, or the respective empty vector controls (pUC19, pBR322 and pJG04), were analyzed using a microplate reader in the absence or presence of the wt λ phage. 200 µl of cultures per well with an OD_600_ of 0.1 in λLB containing ampicillin and either without (A) or with phages corresponding to a MOI of 0.2 (≈ 3.2·10^6^ phage particles) (B), or a MOI of 2.0 (≈ 3.2·10^7^ phage particles) (C) were incubated at 37 °C with agitation (237 cpm) for 15 hours and the OD_600_ and fluorescence were measured every 10 minutes. The protein concentrations were calculated from the fluorescence units using the slopes of the calibration curves (see Figure 4C) and normalized to the background fluorescence of the corresponding empty vector control. Shown are the growth of *E. coli* harboring the depicted plasmids or their corresponding empty vector controls, as well as the changes in MT-mVenus and RE-mCherry concentrations in cultures of *E. coli* harboring the depicted plasmids. (D) Schematic overview of the experiment, illustrating the action of the RE on the DNA of infecting phages. Data represents the mean of three replicates and their standard error (half-transparent shades).

To conclude, expression of the *Cfr*BI R-M system from high-, medium or low-copy plasmids was sufficient to provide effective protection for *E. coli* from phage λ at the investigated concentrations. However, *E. coli* harboring the high-copy empty vector showed increased susceptibility to phages, compared to cells harboring the medium- and low-copy plasmids (Figure 7).

## DISCUSSION

In this study, we investigated the expression dynamics of the DNA-methylation-regulated type II R-M system *Cfr*BI (Figure 1) in *E. coli*, factors influencing the expression of the MT and RE and the ability of the R-M system to protect its host against phages. We show that the R-M system and fluorescent protein fusions are able to protect *E. coli* from bacteriophages (Figure 2 & 3). Next, growth experiments using the fluorescent reporter strains show that expression dynamics in cells harboring the R-M system and in those acquiring it depend on whether a high-, medium-, or low-copy plasmid contains the R-M system (Figure 4 & 5), with the higher the copy number, the higher the overall expression, the higher the ratio of RE to MT and the faster the establishment of the R-M system. In the background of the high-copy plasmid, the overexpression of the MT and RE genes results in a delay of the switch from MT to RE expression after re-inoculating cells already harboring the R-M system. This delay, as well as increased metabolic burden and subsequently cell growth negatively impacts phage resistance. Experiments using a dual-reporter system support the findings by showing that increased expression of the MT represses the MT promoter, while increasing the activity of the RE promoters (Figure 6). Finally, the presence of λ phages does not impact the expression dynamics, but experiments corroborate that cells containing the high copy plasmid pUC19 show decreased phage resistance in comparison to cells harboring medium- or low-copy plasmids (Figure 7).

In contrast to eukaryotes, the role of DNA-modifications in the epigenetic regulation of gene expression in prokaryotes has been less well studied. It is however known to be not only involved in host defense mechanisms but has also been shown to play a role in cell cycle control and the regulation of gene expression of, among others, virulence genes (43, 44). In most bacteria, methylation of the N-6 position of adenine nucleotides (m^6^A) is the predominant form of DNA-methylation, but C-5 cytosine methylation (m^5^C), which is predominantly found in eukaryotes, and N-4 cytosine methylation (m^4^C), like catalyzed by the *Cfr*BI MT, have also been reported in bacteria (45, 46). This m^4^C methylation is less well studied in the context of the regulation of gene expression in bacteria and only in recent years has its role begun to be elucidated in several bacteria (47–51). In past studies, the methylation of the unique *Cfr*BI recognition site in the intergenic region between MT and RE encoding genes, has been shown to interfere with RNA polymerase binding to the stronger MT promoter, facilitating expression of the RE from its weaker promoters upon methylation, shifting the expression from MT to RE (16, 18). However, the dynamics of this regulation and its functional impact have not been studied in detail. The *Cfr*BI R-M system recognizes the relaxed sequence of 5’-CCWWGG-3’ (with W being A or T), which of course has the advantage that it occurs with a higher frequency in genomes and that evading the R-M system by mutations that eliminate the recognition sites is less probable. However, the relaxed recognition site also implies the importance a correct regulation has for the host cell. In the genome of *E. coli* MG1655 (NC_000913) 1,058 recognition sites of the *Cfr*BI R-M systems are present. Incorrect expression of the MT and RE genes, or loss of the MT over time therefore would have devastating consequences for the host cell, resulting in post-segregational killing (52, 53).

Since type II R-M systems, like *Cfr*BI, are often plasmid-based (19), naïve cells that take up such a R-M system containing plasmid require stringent mechanisms that only allow expression of the RE encoding gene upon sufficient expression of the MT. We hypothesize that in a R-M system, in which expression of the genes is only regulated by a single DNA-modification, expression strength and initial gene load of the MT encoding gene determines its expression dynamics, with an initial high ratio of MT to recognition sites favoring faster expression of the RE. Indeed, not only the strength of expression, but dynamics and ratio of RE to MT expression depends on the copy number of the plasmid harboring the system (see Figure 4). Interestingly, in cells already harboring the R-M system, in the background of a low-copy plasmid, expression of the RE gene is constantly lower, indicating that MT expression is not sufficient to keep up with the renewed requirement of re-methylating all of the recognition sites, including the recognition site in the promoter region to allow for more than basal expression of the RE gene. However, it is sufficiently expressed to protect *E. coli* from low to moderate amounts of phages (see Figure 3 and 7). In the background of the medium-copy plasmid, expression of the MT and RE genes was quite similar in strength, however, *E. coli* harboring these plasmids showed the highest resistance to infections by phages (Figure 3). In case of the high-copy plasmids, expression of the MT and RE genes was quite high and the RE accumulated to very high levels during the growth of the bacteria.

Comparing the expression of the MT and RE genes of the *Cfr*BI R-M system R-M systems that have previously been studied, like the C protein-containing *Esp*1396I (54), some similarities but also differences in expression patterns can be noted. In the case of *Esp*1396I, transformation of naïve cells leads to a quick increase in MT expression, followed by a sharp decline and normalization on an intermediate expression level, while expression of the RE increases exponentially after an initial lag phase (26). In the here studied C protein-less R-M system, the initial increase of MT expression is not as steep. However, MT expression also precedes expression of the RE. When RE expression starts to pick up pace, an initial decline in MT expression is observable, but not as pronounced as for *Esp*1396I and MT expression continue to rise afterwards. Since the expression dynamics of the *Cfr*BI R-M system mainly rely on the methylation in the promoter region and the cells are constantly dividing, it fits the expectations that expression of the MT is maintained to a higher extend then in C protein-containing R-M systems in which the concentration of the C protein is the predominant factor. In the case of the *Pvu*II R-M system (55, 56) that also contains a C protein, expression of the RE also lags behind the MT. Expression patterns are similar to expression of the *Cfr*BI R-M system from the low-copy plasmid, since RE expression stays below the levels of MT expression (57). Concerning the impact of MT and RE expression on the protective properties of the R-M system, interesting observations for the *Esp*1396I R-M system were made. Similar to the *Cfr*BI R-M system, an increase in the plasmid copy number results in lower RE to MT ratios of the *Esp*1396I R-M system (32). The study furthermore showed that phage infections during high MT to RE expression favor the occurrence of methylated and therefore resistant phage DNA and that balanced expression of MT and RE are crucial for minimizing this risk for the bacterial population (32). These observations are in good agreement with the phenotypic differences observed for *Cfr*BI (see above): A balanced expression of RE and MT is more important than simply the expression strength of the RE and infection by phages under condition favoring higher MT expression can be detrimental for the bacterial population.

We also wanted to study whether the *Cfr*BI R-M system and its regulation is also functional in the more distantly related bacterium *B. subtilis*, however likely due to different promoter requirements, the MT and RE promoters are not properly expressed (data not shown). Although less well studied than in *E. coli*, DNA methylation has also been shown to regulate gene expression in *B. subtilis* (58, 59) and it would be interesting to further study epigenetic gene regulation in this important model bacterium. It is also interesting to speculate whether the methylation by the *Cfr*BI R-M system impacts the regulation of unrelated genes of the *E. coli* genome. Of the ∼1000 recognition sites, 58 are situated in intergenic regions and may therefore indeed affect the expression of *E. coli* genes and thus have phenotypic consequences. Interestingly, a recent study investigated how horizontal gene transfer of a C protein interferes with the expression of the *racR* gene in *E. coli*, leading to de-repression of a prophage transcriptional regulator and ultimately proves lethal for the host cells, demonstrating how horizontal gene transfer of R-M systems can also be detrimental for their new host (60). Other interesting aspects for future studies are the biochemical properties of the MT and RE of the *Cfr*BI R-M system. While structural predictions indicate that the important residues for *S*-adenosyl methionine binding are conserved (supplementary Figure S1 A), the predicted flexible N-terminus and that the most similar Foldseek hits of the predicted structure are homodimeric could indicate novel binding and DNA-recognition mechanisms of the *Cfr*BI MT. In case of the RE, the predicted catalytic center shows that important residues are present, but that they seem to deviate from the usual motif (supplementary Figure S1 B), also indicating that the *Cfr*BI RE may feature an altered catalytic mechanism interesting for biochemical studies. It is also interesting to speculate how e.g., environmental conditions like nutrient availability, ambient temperature or stress conditions may impact the expression of this epigenetic switch and consequently its protection for the host cells.

To conclude, we demonstrated how different plasmid copy numbers affect the establishment of the *Cfr*BI R-M system, its epigenetic switch of MT to RE expression, the dynamics of their expression and how they and the plasmids impact the ability of the R-M system to protect its host against phages.

## DATA AVAILABILITY

The authors declare that all data supporting the findings of this manuscript are available within the article and its supplementary data files.

## SUPPLEMENTARY DATA

Supplementary Data are available online.

## AUTHOR CONTRIBUTIONS

Conceptualization: J.G., N.E.M., K.S., M.A.K.; Methodology: J.G., N.E.M.; Investigation: J.G., T.D.H., Visualization: J.G.; Writing—original draft: J.G.; Writing—review & editing: J.G., N.E.M., K.S., M.A.K.

## ACKNOWLEDGEMENTS & FUNDING

We are grateful to Lucia Villani and Alexander Kirillov for help with some experiments and to Fabian M. Commichau for laboratory and financial support, as well as critical reading of the manuscript. We thank Dorothee Kiefer for providing λ and T7 phages for some experiments. This work was supported by the Deutsche Forschungsgemeinschaft (DFG, German Research Foundation) within the framework of the Walter Benjamin Programme (project numbers 440930027, 517716052) to J.G.

## CONFLICT OF INTEREST

None declared.

## Notes

### Competing Interest Statement

The authors have declared no competing interest.

## REFERENCES

1. Salmond GP, Fineran PC. A century of the phage: past, present and future. Nat Rev Microbiol. 2015; 13: 777–786.

2. Gordillo Altamirano FL, Barr JJ. Phage therapy in the postantibiotic era. Clin Microbiol Rev. 2019; 32: e00066–18.

3. Jansen R, Embden JD, Gaastra W, Schouls LM. Identification of genes that are associated with DNA repeats in prokaryotes. Mol Microbiol. 2002; 43: 1565–1575.

4. Barrangou R, Fremaux C, Deveau H, et al. CRISPR provides acquired resistance against viruses in prokaryotes. Science. 2007; 315: 1709–1712.

5. Brouns SJ, Jore MM, Lundgren M, et al. Small CRISPR RNAs guide antiviral defense in prokaryotes. Science. 2008; 321: 960–964.

6. Molineux IJ. Host-parasite interactions: recent developments in the genetics of abortive phage infections. New Biol. 1991; 3: 230–236.

7. Swarts DC, Makarova K, Wang Y, et al. The evolutionary journey of Argonaute proteins. Nat Struct Mol Biol. 2014; 21: 743–753.

8. Goldfarb T, Sberro H, Weinstock E, et al. BREX is a novel phage resistance system widespread in microbial genomes. EMBO J. 2015; 34: 169–183.

9. Doron S, Melamed S, Ofir G, et al. Systematic discovery of antiphage defense systems in the microbial pangenome. Science. 2018; 359: eaar4120.

10. Ofir G, Melamed S, Sberro H, et al. DISARM is a widespread bacterial defence system with broad anti-phage activities. Nat Microbiol. 2018; 3: 90–98.

11. Georjon H, Bernheim A. The highly diverse antiphage defence systems of bacteria. Nat Rev Microbiol. 2023; 21: 686–700.

12. Loenen WA, Dryden DT, Raleigh EA, et al. Highlights of the DNA cutters: a short history of the restriction enzymes. Nucleic Acids Res. 2014; 42: 3–19.

13. Kravets AN, Solonin AS, Zakharova MV, Tarutina ZE. Plasmid localization and cloning of restriction modification genes from *Citrobacter freundii* 4111 strain. Mol Gen Mikrobiol Virusol. 1992; 7-8: 4–7.

14. Zakharova MV, Kravetz AN, Beletzkaja IV, et al. Cloning and sequences of the genes encoding the *Cfr*BI restriction-modification system from *Citrobacter freundii*. Gene. 1993; 129: 77–81.

15. Zakharova MV, Beletskaya IV, Ibryashkina EM, Solonin AS. An alternative approach to study the enzymatic specificities of the *Cfr*BI restriction-modification system. Heliyon. 2019; 5: e01846.

16. Beletskaya IV, Zakharova MV, Shlyapnikov MG, et al. DNA methylation at the *Cfr*BI site is involved in expression control in the *Cfr*BI restriction-modification system. Nucleic Acids Res. 2000; 28: 3817–3822.

17. Zakharova MV, Beletskaya IV, Ibryashkina EM, Solonin AS. An alternative approach to study the enzymatic specificities of the *Cfr*BI restriction-modification system. Heliyon. 2019; 5: e01846.

18. Zakharova M, Minakhin L, Solonin A, Severinov K. Regulation of RNA polymerase promoter selectivity by covalent modification of DNA. J Mol Biol. 2004; 335: 103–111.

19. Jeltsch A, Pingoud A. Horizontal gene transfer contributes to the wide distribution and evolution of type II restriction-modification systems. J Mol Evol. 1996; 42: 91–96.

20. Sambrook J, Russell DW. Molecular cloning: a laboratory manual. Cold Spring Harbor Laboratory Press, Cold Spring Harbor, NY.

21. Cohen SN, Chang AC. Revised interpretation of the origin of the pSC101 plasmid. J Bacteriol. 1977; 132: 734–737.

22. Yanisch-Perron C, Vieira J, Messing J. Improved M13 phage cloning vectors and host strains: nucleotide sequences of the M13mp18 and pUC19 vectors. Gene. 1985; 33: 103–119.

23. Gibson DG, Young L, Chuang RY, et al. Enzymatic assembly of DNA molecules up to several hundred kilobases. Nat Methods. 2009; 6: 343–345.

24. Kremers GJ, Goedhart J, van Munster EB, Gadella TW Jr. Cyan and yellow super fluorescent proteins with improved brightness, protein folding, and FRET Förster radius. Biochemistry. 2006; 45: 6570–6580.

25. Shaner NC, Campbell RE, Steinbach PA, et al. Improved monomeric red, orange and yellow fluorescent proteins derived from *Discosoma sp*. red fluorescent protein. Nat Biotechnol. 2004; 22: 1567–1572.

26. Morozova N, Sabantsev A, Bogdanova E, et al. Temporal dynamics of methyltransferase and restriction endonuclease accumulation in individual cells after introducing a restriction-modification system. Nucleic Acids Res. 2016; 44: 790–800.

27. Horton RM, Cai ZL, Ho SN, Pease LR. Gene splicing by overlap extension: tailor-made genes using the polymerase chain reaction. Biotechniques. 1990; 8: 528–535.

28. Waldo GS, Standish BM, Berendzen J, Terwilliger TC. Rapid protein-folding assay using green fluorescent protein. Nat Biotechnol. 1999; 17: 691–695.

29. Merzbacher M, Detsch C, Hillen W, Stülke J. *Mycoplasma pneumoniae* HPr kinase/phosphorylase. Eur J Biochem. 2004; 271: 367–374.

30. Schilling T, Dietrich S, Hoppert M, Hertel R. A CRISPR-Cas9-based toolkit for fast and precise *in vivo* genetic engineering of *Bacillus subtilis* phages. Viruses. 2018; 10: 241.

31. Inoue H, Nojima H, Okayama H. High efficiency transformation of *Escherichia coli* with plasmids. Gene. 1990; 96: 23–28.

32. Kirillov A, Morozova N, Kozlova S, et al. Cells with stochastically increased methyltransferase to restriction endonuclease ratio provide an entry for bacteriophage into protected cell population. Nucleic Acids Res. 2022; 50: 12355–12368.

33. Abramson J, Adler J, Dunger J, et al. Accurate structure prediction of biomolecular interactions with AlphaFold 3. Nature. 2024; 630: 493–500.

34. Meng EC, Goddard TD, Pettersen EF, et al. UCSF ChimeraX: Tools for structure building and analysis. Protein Sci. 2023; 32: e4792.

35. Schneider CA, Rasband WS, Eliceiri KW. NIH Image to ImageJ: 25 years of image analysis. Nat Methods. 2012; 9: 671–675.

36. Guzman LM, Belin D, Carson MJ, Beckwith J. Tight regulation, modulation, and high-level expression by vectors containing the arabinose P_BAD_ promoter. J Bacteriol. 1995; 177: 4121–4130.

37. Bolivar F, Rodriguez RL, Greene PJ, et al. Construction and characterization of new cloning vehicles. II. A multipurpose cloning system. Gene. 1977; 2: 95–113.

38. Lerner CG, Inouye M. Low copy number plasmids for regulated low-level expression of cloned genes in *Escherichia coli* with blue/white insert screening capability. Nucleic Acids Res. 1990; 18: 4631.

39. Lin-Chao S, Chen WT, Wong TT. High copy number of the pUC plasmid results from a Rom/Rop-suppressible point mutation in RNA II. Mol Microbiol. 1992; 6: 3385–3393.

40. Atlung T, Christensen BB, Hansen FG. Role of the rom protein in copy number control of plasmid pBR322 at different growth rates in *Escherichia coli* K-12. Plasmid. 1999; 41: 110–119.

41. Roberts RJ, Vincze T, Posfai J, Macelis D. REBASE: a database for DNA restriction and modification: enzymes, genes and genomes. Nucleic Acids Res. 2023; 51: D629–D630.

42. Mierzejewska K, Bochtler M, Czapinska H. On the role of steric clashes in methylation control of restriction endonuclease activity. Nucleic Acids Res. 2016; 44: 485–495.

43. McGeehan JE, Papapanagiotou I, Streeter SD, Kneale GG. Cooperative binding of the C.*Ahd*I controller protein to the C/R promoter and its role in endonuclease gene expression. J Mol Biol. 2006; 358: 523–531.

44. Seong HJ, Han SW, Sul WJ. Prokaryotic DNA methylation and its functional roles. J Microbiol. 2021; 59: 242–248.

45. Vasu K, Nagaraja V. Diverse functions of restriction-modification systems in addition to cellular defense. Microbiol Mol Biol Rev. 2013; 77: 53–72.

46. Beaulaurier J, Schadt EE, Fang G. Deciphering bacterial epigenomes using modern sequencing technologies. Nat Rev Genet. 2019; 20: 157–172.

47. Nye TM, Jacob KM, Holley EK, et al. DNA methylation from a type I restriction modification system influences gene expression and virulence in *Streptococcus pyogenes*. PLoS Pathog. 2019; 15: e1007841.

48. Kumar S, Karmakar BC, Nagarajan D, et al. N^4^-cytosine DNA methylation regulates transcription and pathogenesis in *Helicobacter pylori*. Nucleic Acids Res. 2018; 46: 3429–3445. Erratum in: *Nucleic Acids Res*. 2018; **46**: 3815.

49. Gaultney RA, Vincent AT, Lorioux C, et al. 4-Methylcytosine DNA modification is critical for global epigenetic regulation and virulence in the human pathogen *Leptospira interrogans*. Nucleic Acids Res. 2020; 48: 12102–12115.

50. Fang JL, Gao WL, Xu WF, et al. m^4^C DNA methylation regulates biosynthesis of daptomycin in *Streptomyces roseosporus* L30. Synth Syst Biotechnol. 2022; 7: 1013–1023.

51. Schmidt N, Stappert N, Nimura-Matsune K, et al. Epigenetic control of tetrapyrrole biosynthesis by m^4^C DNA methylation in a cyanobacterium. DNA Res. 2024; 31: dsae035.

52. Naito T, Kusano K, Kobayashi I. Selfish behavior of restriction-modification systems. Science. 1995; 267: 897–899.

53. Handa N, Kobayashi I. Post-segregational killing by restriction modification gene complexes: observations of individual cell deaths. Biochimie. 1999; 81: 931–938.

54. Cesnaviciene E, Mitkaite G, Stankevicius K, et al. *Esp*1396I restriction-modification system: structural organization and mode of regulation. Nucleic Acids Res. 2003; 31: 743–749.

55. Bart A, Dankert J, van der Ende A. Operator sequences for the regulatory proteins of restriction modification systems. Mol Microbiol. 1999; 31: 1277–1278.

56. Tao T, Bourne JC, Blumenthal RM. A family of regulatory genes associated with type II restriction-modification systems. J Bacteriol. 1991; 173: 1367–1375.

57. Mruk I, Blumenthal RM. Real-time kinetics of restriction-modification gene expression after entry into a new host cell. Nucleic Acids Res. 2008; 36: 2581–2593.

58. Nye TM, van Gijtenbeek LA, Stevens AG, et al. Methyltransferase DnmA is responsible for genome-wide N^6^-methyladenosine modifications at non-palindromic recognition sites in *Bacillus subtilis*. Nucleic Acids Res. 2020; 48: 5332–5348.

59. Xingya Z, Xiaoping F, Jie Z, et al. *Bsu*MI regulates DNA transformation in *Bacillus subtilis* besides the defense system and the constructed strain with *Bsu*MI-absence is applicable as a universal transformation platform for wild-type *Bacillus*. Microb Cell Fact. 2024; 23: 225.

60. Gucwa K, Wons E, Wisniewska A, et al. Lethal perturbation of an *Escherichia coli* regulatory network is triggered by a restriction-modification system’s regulator and can be mitigated by excision of the cryptic prophage Rac. Nucleic Acids Res. 2024; 52: 2942–2960.

