## Supplementary Figures for "Plasmid copy number affects the DNA methylation-driven expression dynamics of the *Cfr*BI restriction-modification system and impacts phage restriction"

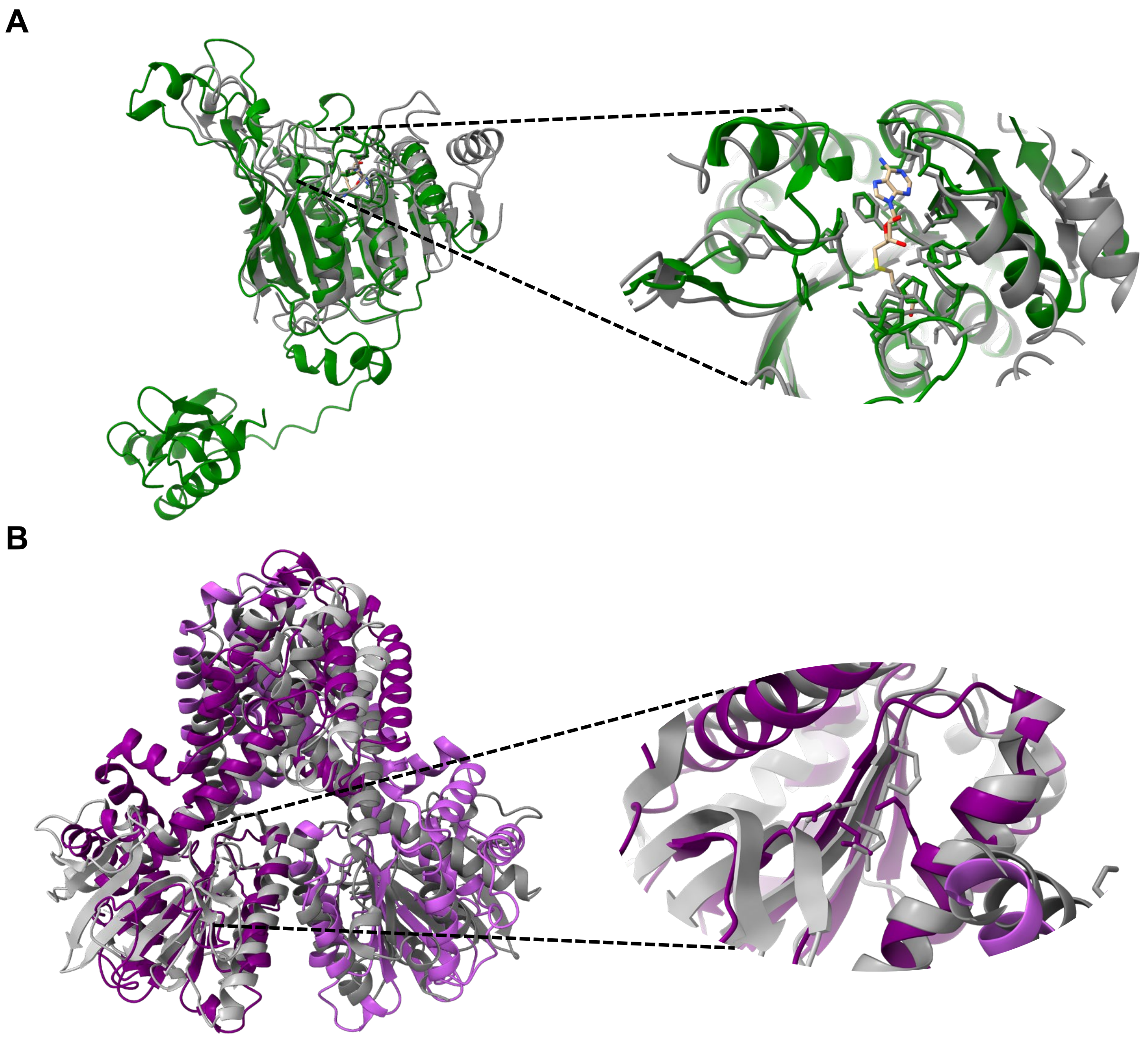


**Supplementary Figure S1. AlphaFold predictions of the *Cfr*BI methyltransferase and restriction endonuclease.** (A) Alignment of the AlphaFold (1) prediction of the *Cfr*BI MT monomer (green) with the crystal structure of the *Pvu*II MT (gray; PDB: 1BOO, (2)). Side chains of amino acids important for *S*-adenosylmethionine-binding are shown in the enlargement. (B) Alignment of the AlphaFold [1] prediction of the *Cfr*BI RE dimer (purple) with the crystal structure of the *Bso*BI RE (gray; PDB: 1DC1, (3)). Side chains of amino acids important for catalysis are shown in the enlargement.

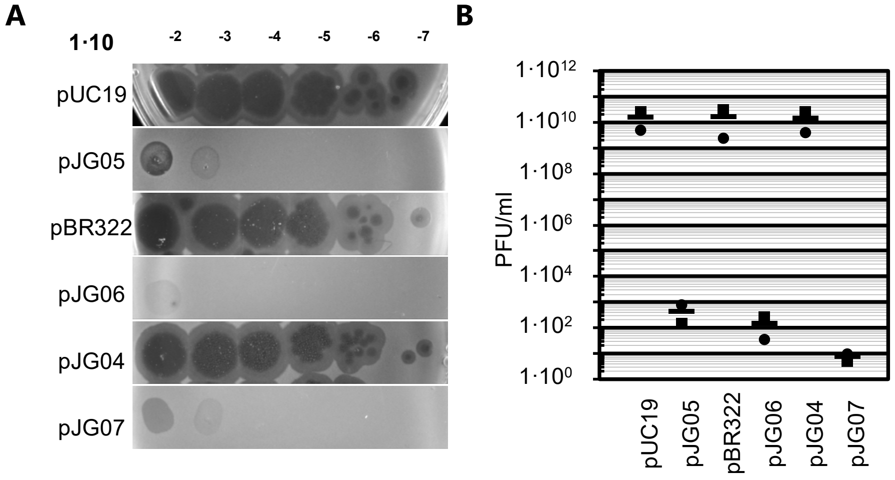


**Supplementary Figure S2**. **The *Cfr*BI restriction-modification system protects *Escherichia coli* against infections by the *E. coli* phage T7.** (A) Phage spot assay to assess phage restriction by the different *Cfr*BI R-M constructs using *E. coli* Top10 strains harboring the indicated plasmids (high copy puC19, medium copy pBR322, low copy pJG04 (pSC101 derivative) or their derivatives harboring the *Cfr*BI R-M system with MT-mVenus and RE-mCherry fusions: pJG05, pJG06 and pJG04, respectively). λLB containing $2{\cdot}10^{10}$ T7 phage particles was 10-fold serial diluted and 5 µl of the indicated dilutions were spotted on agar plates overlaid with λLB top agar, containing the different *E. coli* strains. The plates were photographed after 24 hours of incubation at 37 °C. (B) Graphic representation of the estimated plaque forming units. Plaques of two biological replicates of *E. coli* Top10 strains the indicated plasmids (see (A)) were counted as described in the material and methods and the plaque forming units per ml phage solution was calculated. The bars represent the mean value of the two replicates.

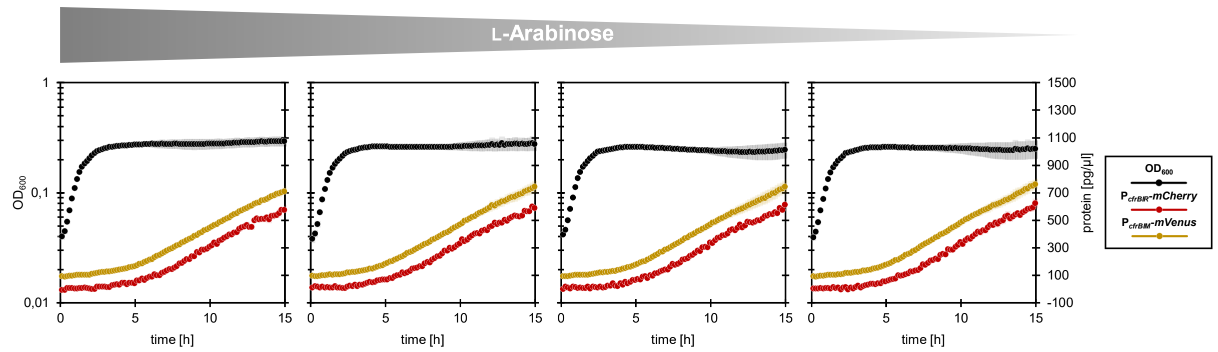


**Supplementary Figure S3**. **Expression of the *Cfr*BI MT is essential to alter the expression dynamics.** *E. coli* Top10 harboring the pBAD33 empty vector and pJG14, a pUC19-derivative that contains $P_{cfrBIM}$-*mVenus* and $P_{cfrBIR}$-*mCherry* fusions were analyzed using a microplate reader. 200 µl of cultures per well with an $OD_{600}$ of 0.1 in LB containing ampicillin, chloramphenicol and different L-arabinose concentrations (from left to right: 0.005, 0.001, 0.0005 and 0 % (w/v)) were incubated at 37 °C with agitation (237 cpm) for 15 hours and the $OD_{600}$ and fluorescence were measured every 10 minutes. The protein concentrations were calculated from the fluorescence units using the slopes of the calibration curves (see Figure 4C) and normalized to the background fluorescence of the corresponding empty vector control. Shown are the growth of *E. coli*, as well as the changes in MT-mVenus and RE-mCherry concentrations.

### References

1. Abramson J, Adler J Dunger J *et al.* . Accurate structure prediction of biomolecular interactions with AlphaFold 3. *Nature*. 2024; **630**: 493-500.
2. Gong W, O'Gara M, Blumenthal RM *et al.* . Structure of *Pvu*II DNA-(cytosine N4) methyltransferase, an example of domain permutation and protein fold assignment. *Nucleic Acids Res*. 1997; **25**: 2702-2715.
3. van der Woerd MJ, Pelletier JJ, Xu S *et al.* . Restriction enzyme *Bso*BI-DNA complex: a tunnel for recognition of degenerate DNA sequences and potential histidine catalysis. *Structure*. 2001; **9**: 133-144.
